# A dual ribosomal system in the zebrafish soma and germline

**DOI:** 10.1101/2024.08.29.610041

**Authors:** Arish N. Shah, Friederike Leesch, Laura Lorenzo-Orts, Lorenz Grundmann, Maria Novatchkova, David Haselbach, Eliezer Calo, Andrea Pauli

## Abstract

Protein synthesis during vertebrate embryogenesis is driven by ribosomes of two distinct origins: maternal ribosomes synthesized during oogenesis and stored in the egg, and somatic ribosomes, produced by the developing embryo after zygotic genome activation (ZGA). In zebrafish, these two ribosome types are expressed from different genomic loci and also differ in their ribosomal RNA (rRNA) sequence. To characterize this dual ribosome system further, we examined the expression patterns of maternal and somatic rRNAs during embryogenesis and in adult tissues. We found that maternal rRNAs are not only expressed during oogenesis but are continuously produced in the zebrafish germline. Proteomic analyses of maternal and somatic ribosomes unveiled differences in core ribosomal protein composition. Most nucleotide differences between maternal and somatic rRNAs are located in the flexible, structurally not resolved expansion segments. Our in vivo data demonstrated that both maternal and somatic ribosomes can be translationally active in the embryo. Using transgenically tagged maternal or somatic ribosome subunits, we experimentally confirm the presence of hybrid 80S ribosomes composed of 40S and 60S subunits from both origins and demonstrate the preferential in vivo association of maternal ribosomes with germline-specific transcripts. Our study identifies a distinct type of ribosomes in the zebrafish germline and thus presents a foundation for future explorations into possible regulatory mechanisms and functional roles of heterogeneous ribosomes.

## Introduction

Ribosomes are macromolecular complexes composed of ribosomal RNAs (rRNAs) and proteins that are responsible for protein synthesis in the cell. Ribosomes of two different origins are present during embryogenesis. Large numbers of maternal ribosomes are synthesized during oogenesis and stored in the egg. Upon fertilization, maternal ribosomes are solely responsible for all translational activity during early embryogenesis (Bazzini et al. 2016; Hensey and Gautier 1997). Maternal ribosomes are progressively degraded during embryogenesis and replaced by newly synthesized ribosomes (i.e., somatic ribosomes) that are produced after zygotic genome activation (ZGA) in the embryo (Heyn et al. 2017; Schramm and Bavister 1999). In addition to their roles in translation (Bazzini et al. 2016; Cenik et al. 2019; Hensey and Gautier 1997; Noack Watt et al. 2016), maternal ribosomes have been proposed to act as a source of pyrimidines required for the synthesis of new ribosomes (Liu et al. 2018b).

In the absence of transcription, the synthesis of new proteins in the early vertebrate embryo needs to be regulated at the post-transcriptional level. Several mechanisms have been described to regulate translation during early embryogenesis, including mRNA localization (Kloc and Etkin 2005; Lorenzo-Orts et al. 2024; Maegawa et al. 1999; Medioni et al. 2012), binding of proteins or miRNAs to mRNAs (Giraldez et al. 2006), and regulation of polyA tail length (Eichhorn et al. 2016; Lim et al. 2016; Liu et al. 2023; Subtelny et al. 2014). While much effort has been devoted to understanding maternal mRNA regulation in the past, we have recently discovered that maternal ribosomes associate with a specific set of factors that bind to key sites on the ribosome and contribute to its repression and stability (Leesch and Lorenzo-Orts et al. 2023). Thus, control of maternal ribosome function also plays an important role in translational regulation during embryogenesis.

Ribosomes have traditionally been considered homogeneous macromolecular complexes with no regulatory role in translation. However, several more recent studies proposed that ribosomes are heterogeneous in their composition (Ferretti et al. 2017; Komili et al. 2007; Mageeney and Ware 2019; O’Leary et al. 2013; Shi et al. 2017; Simsek et al. 2017; Xue et al. 2015). Ribosome compositional heterogeneity is thought to be caused by variations in ribosome accessory factors, ribosome proteins, and ribosomal RNA, as well as by modifications of nucleotides and amino acids comprising the ribosome. For instance, distinct rRNAs have been reported in *Plasmodium berghei* ribosomes isolated from different life cycle stages (Gunderson et al. 1987; Waters et al. 1997), and in ribosomes from *Xenopus* oocytes (Peterson et al. 1980). Moreover, in *Drosophila*, ribosomes isolated from the germline and soma showed a different protein composition (Hopes et al. 2022). Although heterogeneous ribosomes exist in various developmental and cellular contexts, the functional implications of ribosome heterogeneity remain controversial (Barna et al. 2022; Ferretti and Karbstein 2019).

In zebrafish, maternal and somatic ribosomes differ in their composition (Locati et al. 2017a, 2017b), which allows studying these specific ribosome types in an experimentally well-established vertebrate model organism. Maternal ribosomes differ in all four rRNA sequences (28S, 18S, 5.8S, and 5S) and originate from a different ribosomal DNA (rDNA) locus compared to somatic ribosomes (Locati et al. 2017a, 2017b). Maternal rRNAs are transcribed during oogenesis from a single rDNA cluster located on chromosome 4 (Locati et al. 2017b), which is methylated and thus transcriptionally silenced in the soma (Ortega-Recalde et al. 2019). Maternal rRNAs differ from somatic rRNAs in up to 14% of their sequence. However, the consequences of these differences on ribosome functions are not yet known. Furthermore, it remains unclear whether maternal and somatic ribosomes differ in protein composition and are able to form hybrid subunits, which would suggest functional compatibility among their subunits.

Here we analyze the expression, protein composition, and compatibility of maternal and somatic ribosomal subunits in zebrafish. Our results show that despite the differences in rRNA sequences, maternal and somatic ribosomes are found in polysomes indicating their translational activity and can form hybrid ribosomes containing subunits of both origins. Further, we identify germ cells as a cell type that continues to express maternal-type ribosomes throughout development, suggesting a potential functional importance in germ cell-specific translation.

## Results

### Expression of rRNA variants across development and cell types in zebrafish

To investigate the expression of maternal and somatic rRNA variants during zebrafish development, we analyzed total RNA during the first five days of embryogenesis. Consistent with previous findings (Locati et al. 2017a, 2017b), we observed that maternal rRNAs originating from the maternal rDNA locus on chromosome 4 (GRCz11; 4: 77,564,140 – 77,555,053) were the sole rRNA variants present in the egg and during early stages of embryogenesis before zygotic genome activation (ZGA). Following ZGA, somatic rRNAs started to be transcribed from the somatic rDNA locus on chromosome– 5 (GRCz11; 5: 827,807 – 819,029). The levels of somatic rRNAs increased progressively and reached the same level as maternal rRNAs between 20 to 30 hours post-fertilization (hpf). By 5 days post-fertilization (dpf), more than 98% of the detected rRNAs were of somatic origin, reflecting the transition from maternal to somatic ribosomes (**Fig. 1A** and **1B**).

**Figure 1:**
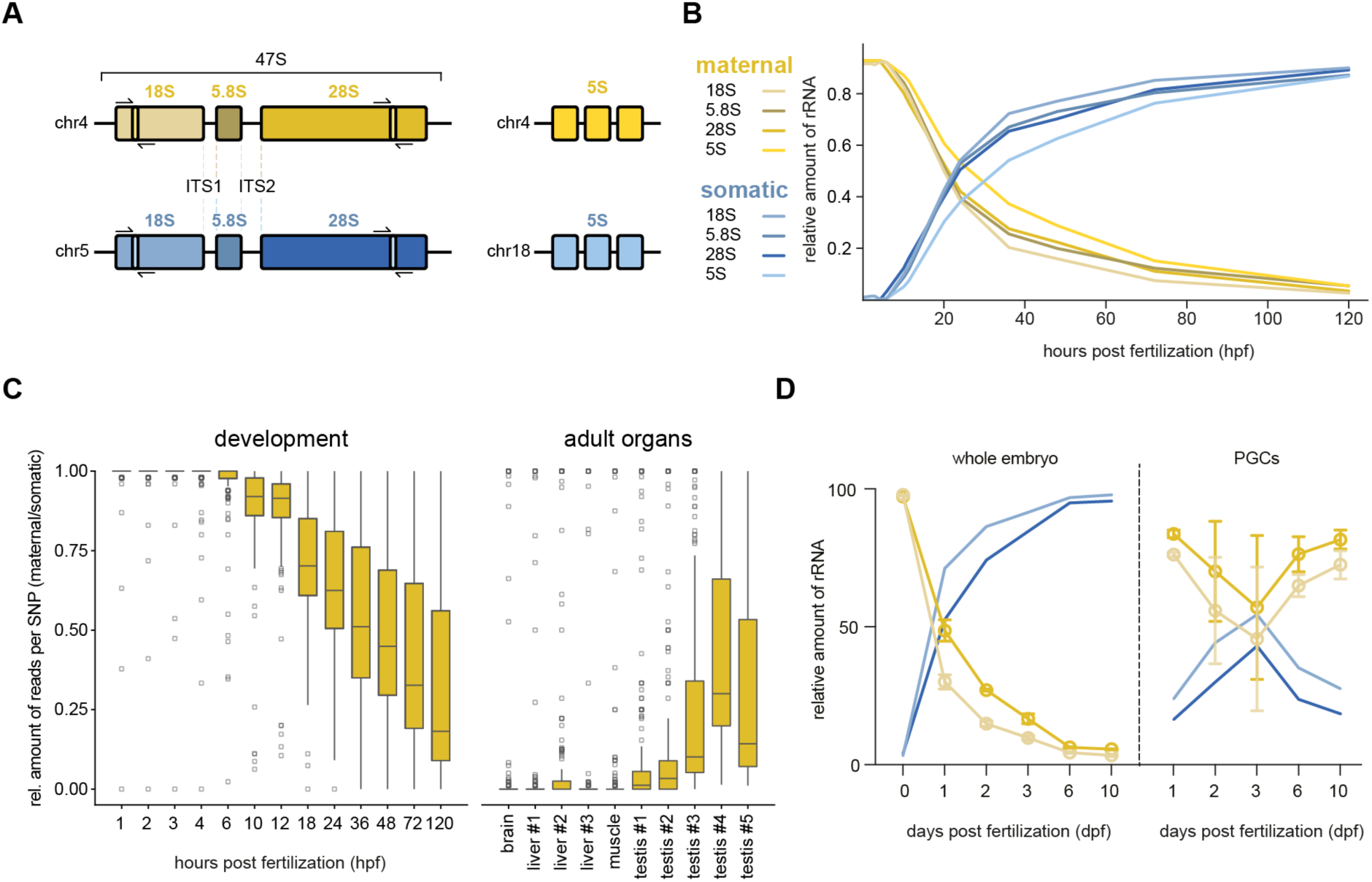
Expression of maternal and somatic rRNA variants in zebrafish. **A)** Organization of maternal (yellow) and somatic (blue) rDNA genes in the zebrafish genome. The 5S rRNA genes are encoded separately, while 18S, 5.8S, and 28S are derived from a single 47S pre-rRNA transcript containing additional spacer sequences (ITS1 and ITS2 (internal transcribed spacers)) that are removed in several processing steps. Expansion segments (ES3S and ES31L) are indicated by lighter-colored boxes and PCR primer icons (see **Supplementary Fig. S1A**). **B)** Relative expression levels of the four rRNAs (28S, 5.8S, 18S, and 5S), comparing maternal (yellow) and somatic (blue) variants. **C)** Fraction of maternal versus somatic rRNAs, obtained by quantifying reads with variant-specific SNPs (single nucleotide polymorphisms) during the first five days of development, in adult somatic organs, and in testes of 5 different males. **D)** Expression of maternal and somatic rRNA variants in whole embryo/larva lysate and FACS-sorted PGCs. ES, expansion segment. FACS, fluorescence activated cell sorting.

Although most ribosomes are of somatic origin by day 5 of development, maternal rRNA variants may be expressed in specific cell types or tissues beyond oocytes. To investigate this, we employed a PCR-based assay that could discriminate between maternal and somatic rRNA variants based on the amplification of highly variable Expansion Segments (ESs) in 18S and 28S RNAs (**Supplementary Fig. S1A**). To confirm the applicability of the PCR-based assay, we recapitulated the transition from maternal to somatic 18S rRNA (**Supplementary Fig. S1B** and **S1C**). While in most adult tissues a single DNA band corresponding to somatic 18S and 28S rRNAs was detected, testis samples showed evidence for the presence of both somatic and maternal rRNAs (**Supplementary Fig. S2**). Next-generation sequencing (NGS) experiments using total RNA from testes dissected from five different adult male zebrafish revealed variable levels of maternal and somatic rRNAs, with maternal rRNA variants accounting for 2% to 27% of the total rRNAs (**Fig. 1C**). Together, these results suggest that both maternal and somatic ribosomes are present in the male germline.

To gain deeper insights into the expression of maternal rRNAs in the germline, we re-analyzed total RNA sequencing data from whole embryos and larvae, as well as published total RNA sequencing data from FACS-sorted primordial germ cells (PGCs) during larval development up to 10 dpf (Redl et al. 2021). PGCs, which are the precursors of the adult germline, showed persistently high levels of maternal rRNAs compared to somatic larval tissues throughout 10 dpf (**Fig. 1D**). Internal transcribed spacer (ITS) sequences are removed from the nascent 47S rRNA and act as a marker for ribosome biogenesis. Consistent with a potential role for maternal ribosomes in the germline, expression analysis of maternal ITS1 (**Fig. 1A**), indicated active transcription of maternal rRNAs in PGCs, but not in the soma, starting from 3 dpf (**Supplementary Fig. S3**); long before the production of early-stage oocytes between 10-25 dpf in juveniles (Dranow et al. 2016; Takahashi 1977).

Despite our evidence for persistent expression of the rDNA locus on chromosome 4 in the zebrafish germline, for simplicity and consistency with previous publications, we will continue to refer to the rRNAs, the subunits they compose, and the ribosomes generated from this locus as “maternal” throughout this manuscript.

Recent interrogation of human cancer biopsies in The Cancer Genome Atlas (TCGA) (Liu et al. 2018a) has similarly shown differential expression of rRNA variants (Rothschild et al. 2023). To test whether zebrafish maternal-type rRNAs might also be re-expressed in immortalized cells or during tumorigenesis, we analyzed cDNAs from several cultured zebrafish cell lines and excised adult tumors (Berghmans et al. 2005; Driever and Rangini 1993; Heilmann et al. 2015; Patton et al. 2005; Paw and Zon 1999; Perez et al. 2018). 18S rRNA compositional analysis in the assayed *ex vivo* or neoplastic contexts showed no evidence of reactivated maternal rRNA (**Supplementary Fig. S4A** and **S4B**). Similarly testing regenerating fin blastema through 5 days post amputation indicates dedifferentiated cells losing morphology and re-entering the cell cycle (Pfefferli and Jaźwińska, 2015) do not express maternal rRNA (**Supplementary Fig. S4C**).

### Maternal and somatic ribosomes differ in protein composition

Maternal and somatic ribosomes have so far only been shown to differ in their rRNAs. To investigate potential differences in protein composition, we performed proteomic analyses of ribosomes isolated at different times during embryogenesis and compared the expression of core ribosomal proteins (RPs) (**Supplementary Table S1**). We chose time points containing either only maternal ribosomes (egg, 1 hpf, 3 hpf and 6 hpf) (Leesch and Lorenzo-Orts et al., 2023) or only somatic ribosomes (120 hpf) (**Fig. 1B**). We detected 8 RPs with more than one paralog associated to maternal or somatic ribosomes during embryogenesis, namely Rps8 (eS8), Rps17 (eS17), Rps26 (eS26), Rps27 (eS27), Rplp2 (P2), Rpl5 (uL18), Rpl7 (uL30) and Rpl22 (eL22) (**Fig. 2A**). Moreover, alternative isoforms were evident for Rps5 (uS7), Rps18 (uS13), Rps19 (eS19) and Rpl9 (uL6) (**Fig. 2A**).

**Figure 2:**
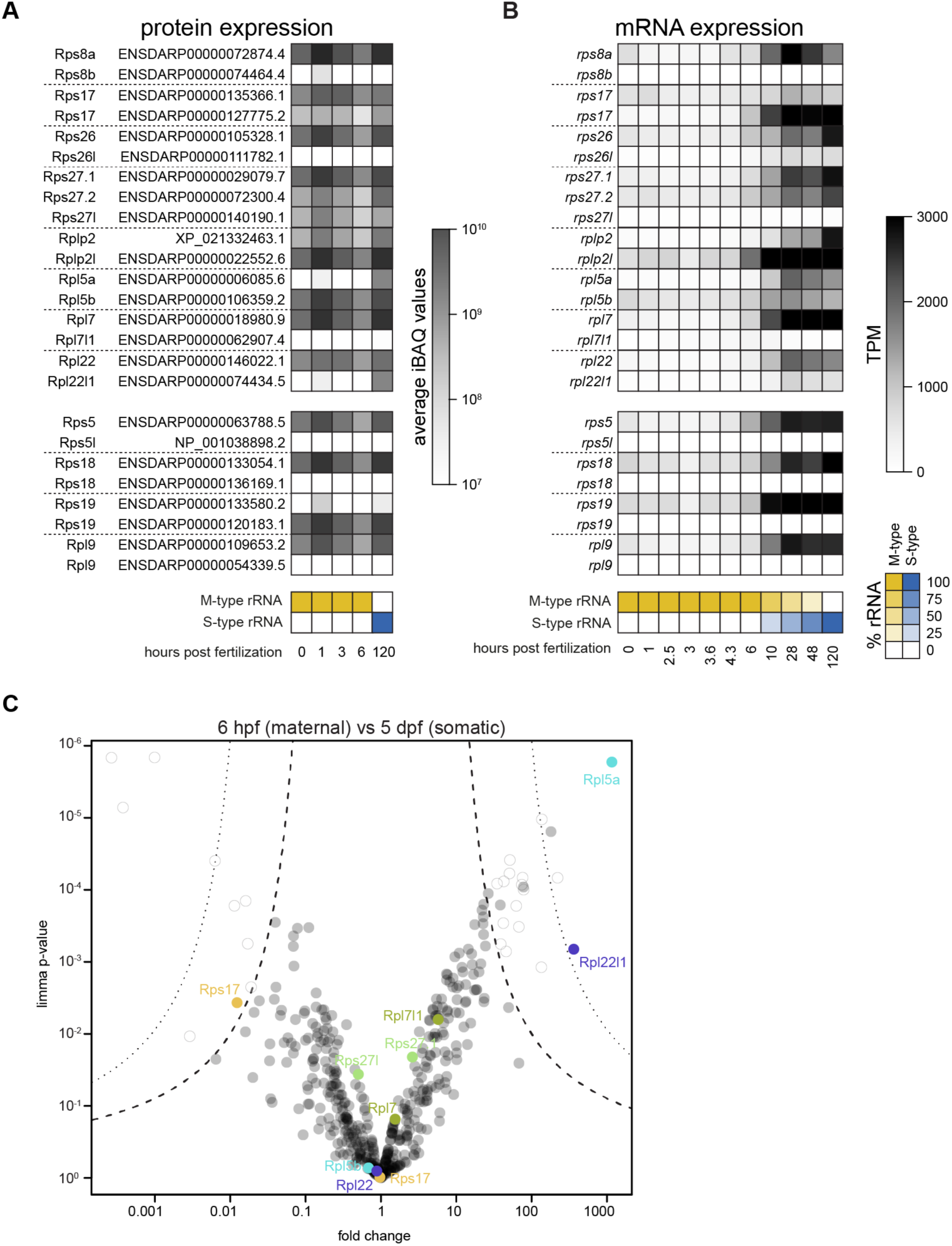
Expression of alternative ribosomal core proteins in zebrafish development. **A)** Heatmap of protein expression (normalized IBAQ values) for variant ribosomal protein pairs (paralogs, top; alternative isoforms, bottom) associated with zebrafish ribosomes at different developmental stages. Below, relative levels of maternal (yellow) and somatic (blue) rRNA variants at these developmental stages are indicated separately. **B)** Heatmap of mRNA expression of paralogs and alternative isoforms for ribosomal protein variants during embryogenesis. See also **Supplementary Fig. S5**. The relative levels of maternal (yellow) and somatic (blue) rRNA variants at these developmental stages are indicated in the scheme below the heat map. **C)** Volcano plot based on mass spectrometry data showing fold enrichment of proteins in the ribosome fraction of 5 dpf larvae versus 6 hpf embryos (n = 3 for each time point). Pairs of ribosomal protein variants with significantly different associations are shown in different shades. Non-significant but detected variants are shown in green. Permutation-based false discovery rates (FDRs) are shown as dotted (FDR < 0.01) and dashed (FDR < 0.05) lines. All significantly enriched or depleted proteins are listed in **Supplementary Table S1**. TPM, transcripts per million. iBAQ, intensity Based Absolute Quantification.

To gain further insights into the expression patterns of these alternative RPs, we analyzed previously published RNA-seq datasets covering different stages of oogenesis, embryogenesis, and adult tissues (Cabrera-Quio et al. 2021; Noda et al. 2022; Pauli et al. 2014) (**Fig. 2B, Supplementary Fig. S5**). While only one protein variant was predominantly present (more than 10-fold difference in expression level) for all RPs (**Fig. 2A**), mRNAs encoding for Rps17, Rps27.1/Rps27.2, Rplp2/Rplp2l and Rpl5a/Rpl5b were expressed at more similar levels during development and in adult tissues (**Fig. 2B, Supplementary Fig. S5**). In some cases, one RP paralog was associated with purified ribosomes at all stages while the other was ribosome-bound only at specific times. Examples include Rpl5b and Rpl22, which were associated with ribosomes throughout embryogenesis, whereas their paralogs Rpl5a and Rpl22l1, respectively, only showed increased ribosome incorporation at 5 dpf, correlating with the expression of somatic rRNA variants (**Fig. 2B** and **2C**). This proteomic analysis of maternal and somatic ribosomes during embryogenesis provided evidence for RP composition differences in assembled ribosomes at different times of development, but did not reveal a single RP paralog pair that mirrored the switch between maternal and somatic rRNAs.

### Nucleotide differences between maternal and somatic rRNAs are predominantly located in structurally non-resolved flexible areas of the ribosome

rRNAs provide the ribosome with a structural framework and form many of its core functional sites. Since maternal and somatic ribosomes contain different rRNAs and RPs, we mapped the sites of nucleotide differences onto the previously obtained cryo-EM structure of the maternal ribosome at 6 hpf (Leesch and Lorenzo-Orts et al. 2023) (**Fig. 3A, Supplementary Fig. S6A**). We chose the 6 hpf time-point since it contains exclusively maternal ribosomes (and thus maternal rRNAs) that have already lost the repressive dormancy factors that are initially bound at 1 hpf, and embryos already contain a larger amount of polysomes than at 1 hpf (Leesch and Lorenzo-Orts et al. 2023),

**Figure 3:**
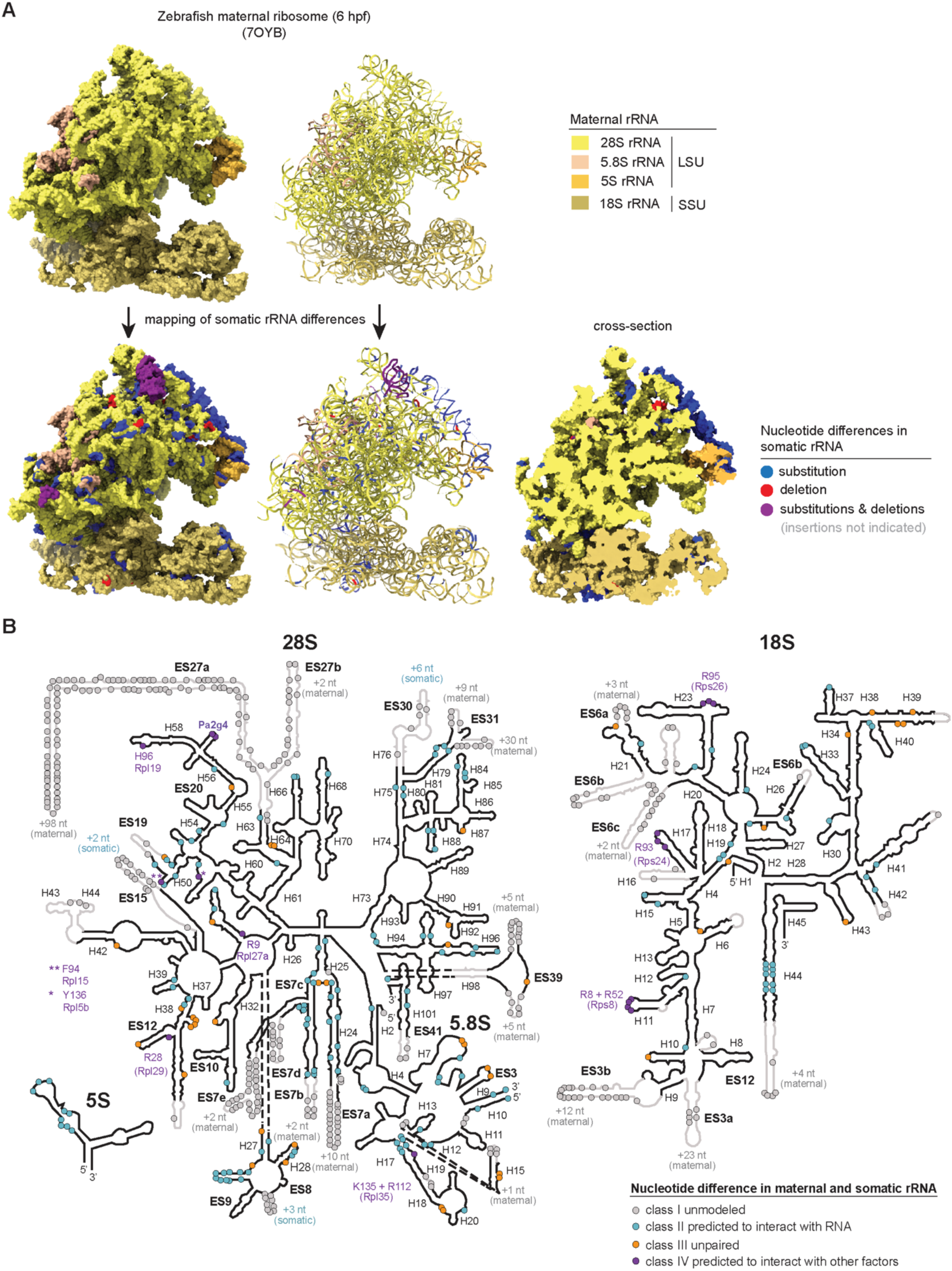
Location of sequence and structural differences in rRNA variants. **A)** Map of the maternal zebrafish ribosome isolated from 6 hpf embryos (PDB-7OYB; Leesch and Lorenzo-Orts et al., 2023). Maternal rRNA sequences are shown in surface representation (left) and cartoon representation (right) in shades of yellow. Locations of nucleotide differences in zebrafish somatic rRNAs are highlighted at the bottom (see legend for color code). A cross-section through the model is shown in the right, revealing that most of the sequence differences are located on the surface and not in the inside of the ribosome’s core. **B)** Sequence differences in maternal and somatic rRNA variants mapped onto rRNA secondary structure predictions derived from R2DT (Sweeney et al., 2021). 18S rRNA, 5.8S and 5S: structural predictions for *Danio rerio* somatic rRNAs; 28S rRNA: structural prediction for the *Homo sapiens* 28S rRNA, with the ES regions adjusted based on modelling of the zebrafish sequence in the Vienna RNAfold package (Lorenz et al., 2011). Circles indicate positions of nucleotide differences between the maternal and somatic rRNAs. Regions not modeled in the cryo-EM structure obtained from 6 hpf zebrafish embryos due to poor density are shown in gray.

As previously suggested (Locati et al. 2017a) the majority of sequence differences between maternal and somatic rRNAs are located in expansion segments (ESs), which are flexible RNA elements on the surface of the ribosome and thus often not resolved by structural methods (**Fig. 3A** and **3B, Movies 1-3**). In fact, only 27% of the nucleotides that differ between maternal and somatic rRNAs were visible in our maternal ribosome structure. In several instances, complementary nucleotide differences were predicted to be present in Watson-Crick base-pairs (e.g. H17, H44, ES27; **Fig. 3B**), thus likely preserving rRNA interactions and secondary structure (**Supplementary Table S2**).

Functionally important nucleotides tend to be conserved throughout evolution. We therefore also analyzed whether the differences between maternal and somatic rRNAs corresponded to evolutionarily conserved or more variable nucleotides. We found that nucleotides that differed between zebrafish maternal and somatic rRNAs were depleted from conserved regions (characterized by low Shannon entropies; Schmitt and Herzel 1997) and instead enriched in areas with high degrees of variability at the single nucleotide level across evolution (high Shannon entropies) (**Supplementary Fig. S6B** and **S6C**). Together, our analyses reveal that the majority of nucleotide differences between maternal and somatic ribosomes are located in evolutionary less conserved regions outside of the structurally well-resolved areas of the ribosome.

### Formation of hybrid ribosomes comprising maternal and somatic subunits

To investigate whether maternal and somatic ribosomal subunits may interact and form 80S “hybrid” ribosomes, we focused our structural analyses on rRNA sequence differences and alternative ribosomal proteins in the contact regions between 40S and 60S subunits, termed intersubunit bridges (see **Table 1**). 40S and 60S subunits interact through 17 bridges (B1a-eB14) (Anger et al. 2013; Ben-Shem et al. 2011; Melnikov et al. 2012). Most of these bridges are formed by RNA-RNA or RNA-protein contacts, with the exception of B1b/c, which is formed by the contact of two ribosomal proteins, Rps18 (uS13) and Rpl11 (uL5). Five bridges are specific for eukaryotic ribosomes (eB8, eB11, eB12, eB13, and eB14). On the RNA level, there are sequence differences between maternal and somatic rRNAs in intersubunit bridges B3-B6, eB13, and in expansion segments ES31L, ES41L, and ES6S (**Fig. 3B**), which form bridges eB8, eB11, eB12 and eB14, respectively (**Table 1**). In addition to rRNA sequence differences, one ribosomal protein, Rps8 (eS8), known to form an intersubunit bridge (Tamm et al. 2019) has a paralogous RP and both are expressed in early zebrafish embryogenesis (**Fig. 2A**). However, in our data, Rps8a was more abundant at the protein and mRNA level in both maternal and somatic ribosomes, suggesting no difference in RP paralog usage at the intersubunit bridge between maternal and somatic ribosomes (**Fig. 2A** and **2B**).

**Table 1:**
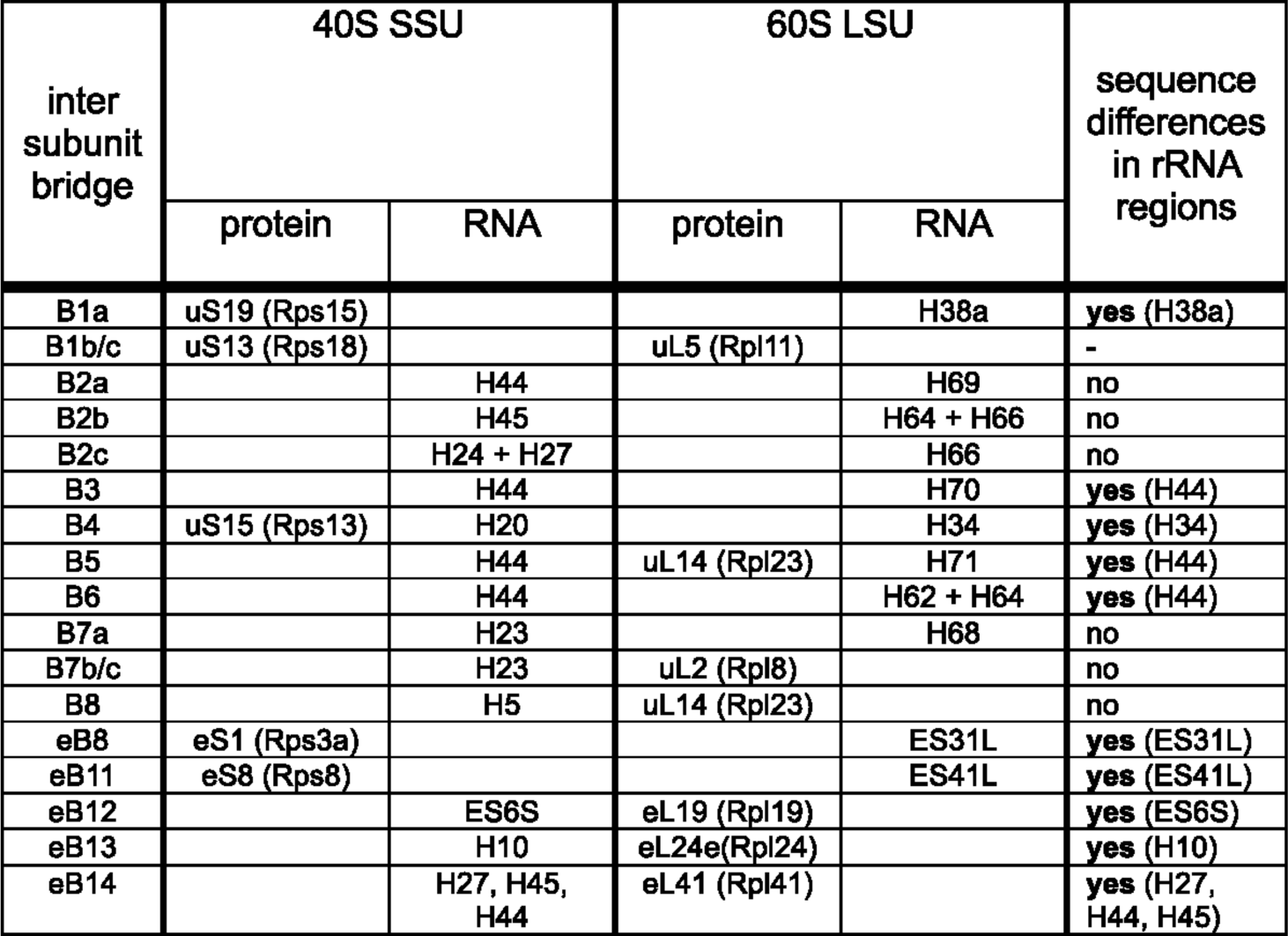
Structural differences of intersubunit bridge regions in maternal and somatic ribosomes. Intersubunit bridges (listed in the first column) are connections between 40S SSU (composed of the 18S rRNA and 33 RPs) and 60S LSU (composed of the 5S, 5.8S, 28S rRNAs, and 46 RPs). Bridges composed of rRNA sequences with differences between maternal and somatic types are marked. H, helix. ES, expansion segment.

To directly investigate the possibility of hybrid ribosome formation *in vivo*, 24 hpf zebrafish larvae were used since maternal and somatic rRNA variants are present at equal levels at this developmental stage (**Fig. 1B, Supplementary Fig. S1B**). To assess whether maternal ribosomes can be found in polysomes at this developmental stage, we performed polysome gradients of total lysates, which showed peaks for the 40S and 60S ribosomal subunits, a large 80S (monosome) peak and several polysome peaks (**Fig. 4A**). Besides detecting maternal and somatic rRNA types in the fraction corresponding to 40S, 60S and 80S subunits, we identified both rRNA types in translationally active polysomes (**Fig. 4B**; see **Supplementary Fig. S1B** for the specificity of the primers), suggesting that both maternal and somatic ribosomes support translation at this stage of embryogenesis.

**Figure 4:**
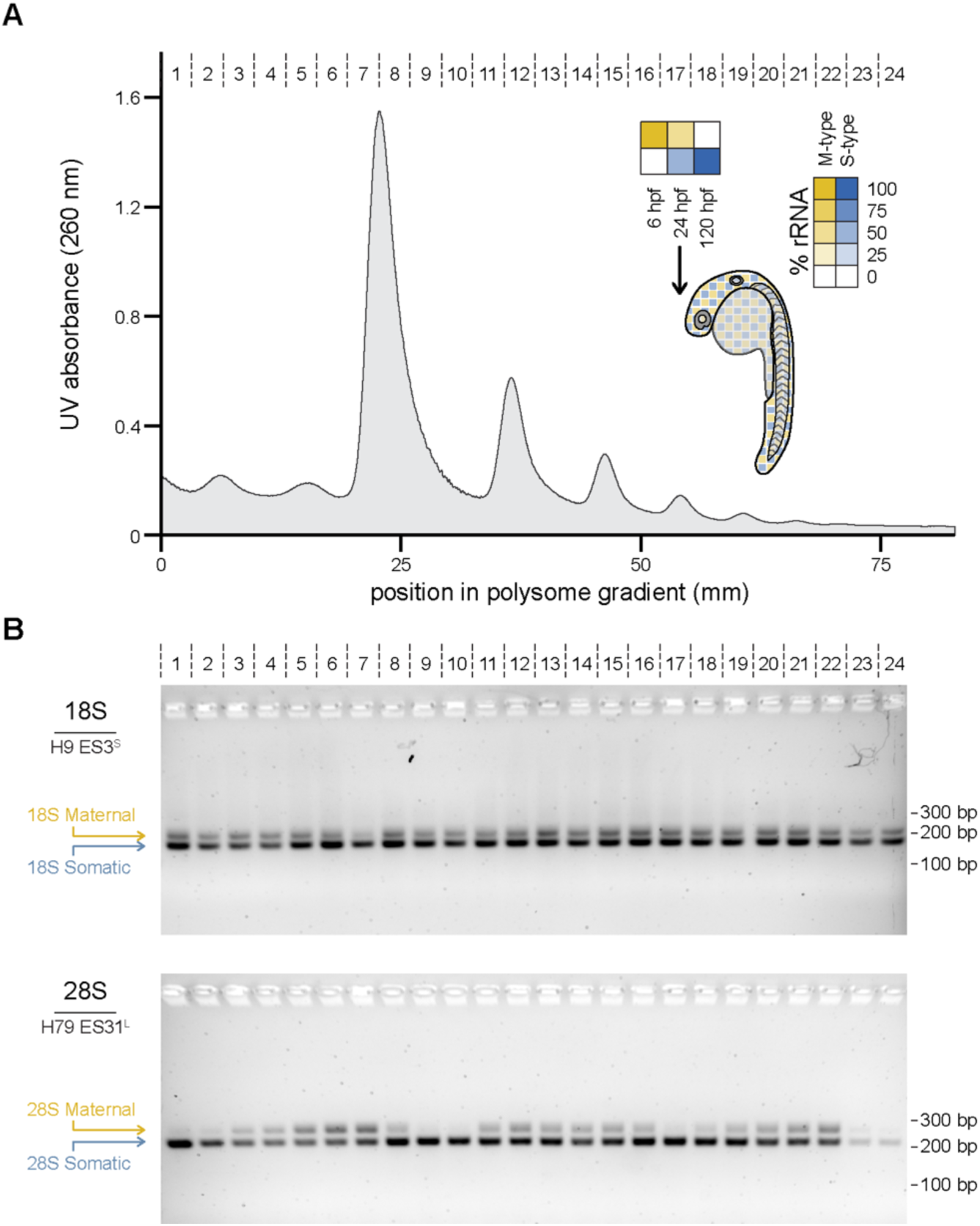
Maternal and somatic ribosomal subunits are translationally active in 24 hpf embryos. **A)** Representative polysome profile with continuous UV absorbance (A260) reading from sucrose density gradients containing lysate from 24 hpf zebrafish embryos for which the relative levels of each rRNA variant is shown in the inset. Position in the gradient and collected fractions are indicated. **B)** Gel electrophoretic analysis of PCR-based detection of maternal and somatic small (18S) and large (28S) rRNA variants in the individual factions indicated in (A). See **Supplementary Fig. S1B** for the specificity of the primers and **Supplementary Table S3** for lengths of PCR products from maternal and somatic rRNAs. bp (base pair), H (helix), ES (expansion segment).

To achieve selective labeling of maternal and somatic ribosomal subunits, FLAG-tagged Rpl10a was specifically expressed during oogenesis via the oogenesis-specific promoter *sycp* (synaptonemal complex protein) (Tg(Mat-RiboFLAG), ensuring the restricted incorporation into maternal ribosomes. In contrast, a male-contributed transgene using a ubiquitously expressed promoter *ubi* (ubiquitin b) (Tg(Som-RiboFLAG)) enabled specific labeling of somatic ribosomes (**Supplementary Fig. S7, S8** and **S9**).

To validate our experimental setup (**Fig. 5A**), we first performed FLAG-immunoprecipitation (FLAG-IP) experiments using maternal and somatic FLAG-tagged Rpl10a embryo lysates treated with EDTA to dissociate 80S ribosomes into individual subunits. By analyzing the relative abundance of maternal and somatic 28S rRNA in the input and IP fractions using RT-qPCR, we confirmed the specificity of our tagging, precipitation, and detection approaches (**Fig. 5B**).

**Figure 5:**
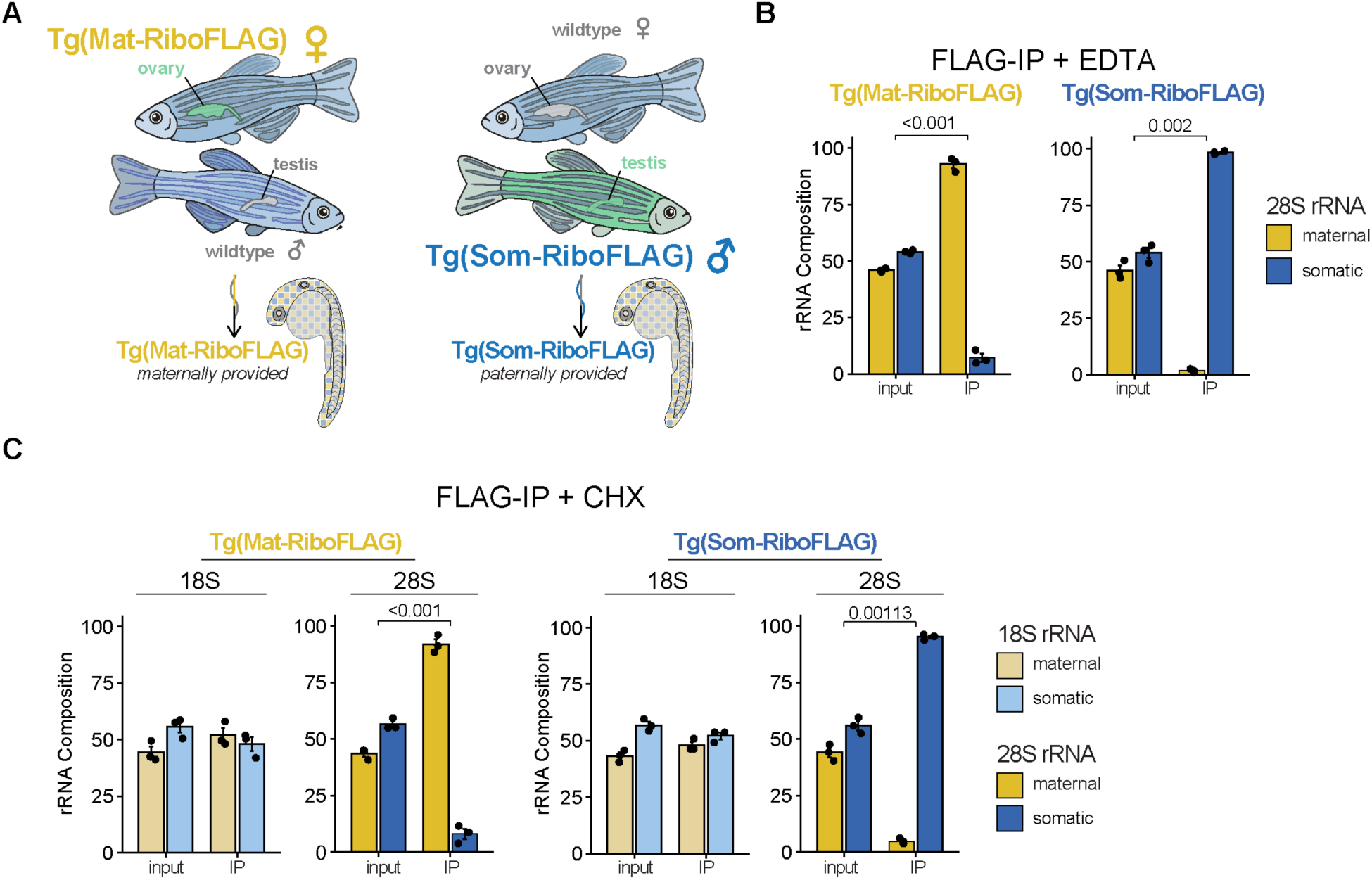
In vivo evidence for hybrid ribosome formation in 1 day post-fertilization embryos. **A)** Scheme summarizing the generation of embryos with tagged ribosomes by crossing Tg(Mat-RiboFLAG) females to wildtype males and Tg(Som-RiboFLAG) males to wildtype females (see Methods). **B)** RT-qPCR analysis of the relative amounts of maternal and somatic 28S rRNAs detected in input or eluate (IP) fractions of a FLAG-IP experiment, which used EDTA-treated lysates containing either tagged maternal (Tg(Mat-RiboFLAG)) or somatic (Tg(Som-RiboFLAG)) 60S subunits as input. **C)** RT-qPCR analysis of the relative amounts of maternal and somatic 28S and 18S rRNAs detected in input or eluate (IP) fractions of a FLAG-IP experiment. Cycloheximide (CHX) was added during lysis of embryos containing either tagged maternal (Tg(Mat-RiboFLAG)) or somatic (Tg(Som-RiboFLAG)) 60S subunits. Monosome fractions from RNAse-treated lysates were used as input. In (B) and (C), three biological replicates from independent natural crosses were used, and significance was calculated by t-test (if not indicated, p-value > 0.05).

To test whether hybrid ribosomes form *in vivo*, we performed a second FLAG-IP experiment in the presence of cycloheximide (CHX), a well-established translation inhibitor known to stabilize translating ribosomes. This allowed us to assess the presence of maternal or somatic small subunits (SSUs) in the IP fraction of the tagged maternal or somatic LSUs (**Fig. 5C**). Intriguingly, we observed similar levels of maternal and somatic SSUs in the IP fractions independently of whether maternal or somatic tagged LSUs had served as baits. These findings provide direct experimental evidence for the *in vivo* formation of hybrid ribosomes, supporting the compatibility between large and small ribosomal subunits of different origins.

### Co-enrichment of maternal ribosomes with PGCs and PGC-specific mRNAs

Building on our observation that maternal rRNAs are continuously expressed in PGCs (**Fig. 1D, Supplementary Fig. S3B**) and that maternal ribosomes are actively translating in 1-day-old zebrafish embryos, we reasoned that maternal ribosomes should be present in PGCs and may be translating PGC-specific transcripts. To test these hypotheses, we injected *in vitro* transcribed *eGFP* mRNA fused to the *nanos3-3’UTR* into the 1-cell stage embryo. The *nanos3-3’UTR* stabilizes the transcript only in PGCs, enabling PGC-specific eGFP translation at 1 dpf (**Fig. 6A** and **6B**).

**Figure 6:**
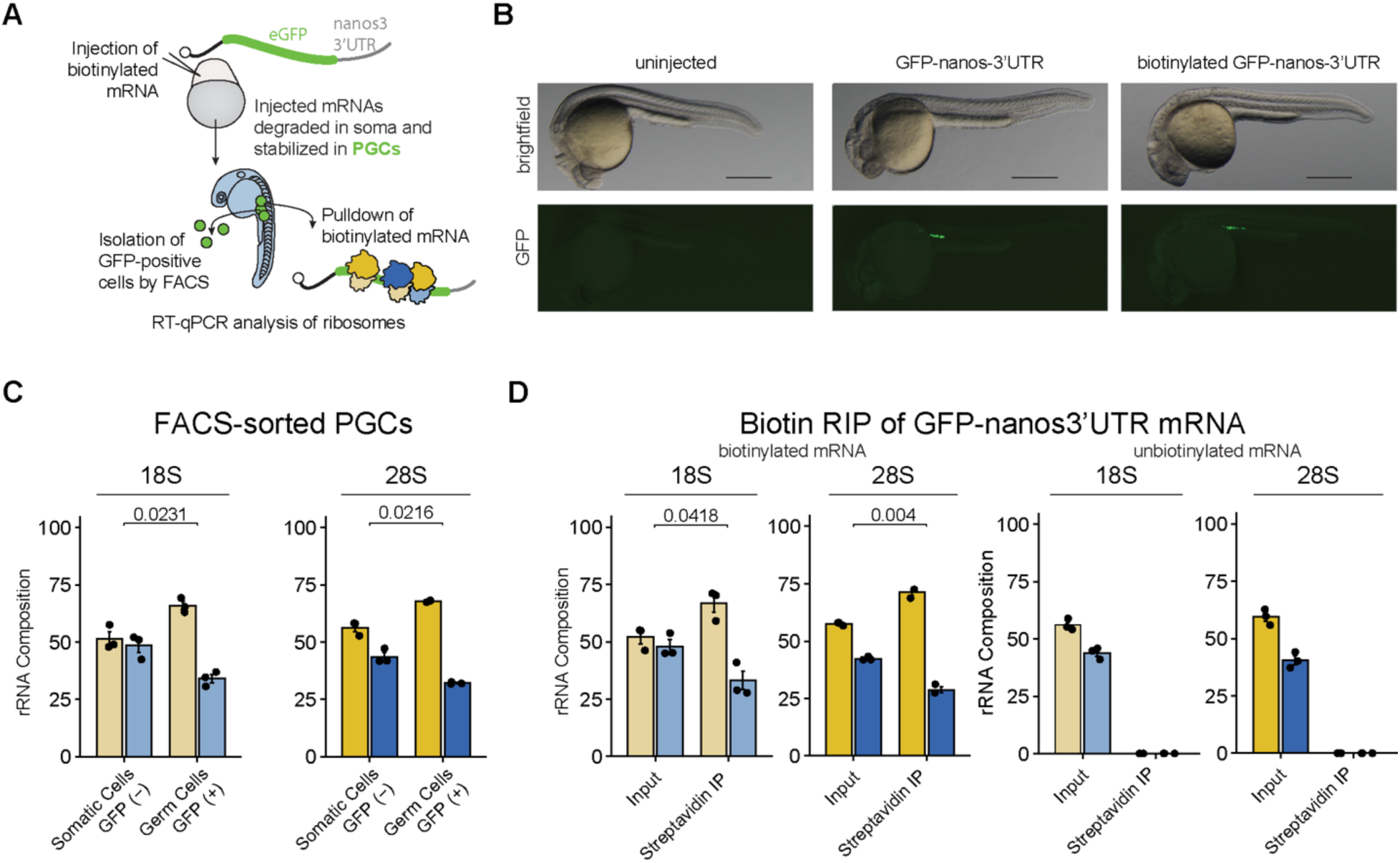
Enrichment of PGC-localized mRNAs with maternal ribosomes at 1 day post-fertilization. **A)** Schematic of the experimental strategy. eGFP-nanos3’UTR containing mRNA was injected into 1-cell stage zebrafish embryos from which two different experimental approaches were used to investigate the ratio of maternal and somatic ribosomes in PGCs and to test which ribosomes are bound to PGC-localized mRNAs. **B)** Brightfield and eGFP images of 1 day post-fertilization embryos, previously injected with only water, eGFP-nanos3’UTR mRNA, or biotinylated eGFP-nanos3-3’UTR mRNA. Scale bars indicate 500 μm. **C)** Ratio of maternal (yellow) versus somatic (blue) 18S and 28S rRNA in FACS-isolated eGFP-positive (PGCs) and eGFP-negative (somatic) cells analyzed by RT-qPCR. **D)** Ratio of maternal (yellow) versus somatic (blue) 18S and 28S variants associated with biotinylated eGFP-nanos3-3’UTR mRNAs isolated by RIP (RNA immunoprecipitation). Non-biotinylated eGFP-nanos3-3’UTR mRNA served as control. Non-specific (background) amplification was determined using IPs from wild-type (uninjected) lysate and was set to zero. In (C) and (D), three biological replicates from independent natural crosses were used, and significance was calculated by t-test (if not indicated, p-value > 0.05).

To confirm that PGCs maintain a higher ratio of maternal versus somatic ribosomes at 1 dpf (**Fig. 1D**), we used RT-qPCR to assess the ratio of maternal and somatic rRNA variants in FACS-isolated eGFP-positive PGCs and non-eGFP expressing somatic cells. As expected, a higher abundance of maternal 18S and 28S rRNA variants were detected in the isolated PGCs compared to somatic cells, confirming the persistence of maternal ribosomes at higher levels in germline cells (**Fig. 6C**). To directly test whether maternal ribosomes associate with PGC-specific transcripts and thus may contribute to germ cell-specific translation, we performed a RNA immunoprecipitation (RIP) assay using biotinylated *eGFP-nanos3-3’UTR* mRNAs extracted from 1-day-old cycloheximide-treated larvae. This approach allowed us to analyze the pool of ribosomes bound to this specific PGC-localized mRNA. Notably, we observed a significant enrichment of the maternal ribosomes compared to somatic ribosomes associated with biotin-labeled PGC-localized mRNAs (**Fig. 6D**). However, this apparent level of enrichment was similar to the level of maternal ribosomes in isolated PGCs (**Fig. 6C**), providing no evidence towards selectivity of maternal ribosomes for germ cell-specific transcripts. Nevertheless, our results reveal an association between maternal ribosomes and PGCs, and thus germline-specific transcripts.

## Discussion

Previous studies have revealed that zebrafish ribosomes of maternal and somatic origin are composed of two distinct sets of rRNAs (Locati et al. 2017a, 2017b). While one set of rRNAs was thought to be exclusively present in maternal ribosomes, our PCR-based and RNA-seq analyses indicate that these rRNAs are not only transcribed during oogenesis, but are also present in PGCs and the male germ line. Translation in PGCs is tightly regulated and differs from that in the soma, as these cells are specified before the onset of zygotic genome activation (ZGA) and contain specific maternal mRNAs (Dranow et al. 2016; Johnson et al. 2011; Yu et al. 2016). Through at least 10 dpf, both inherited and zygotically generated maternal ribosomes constitute the major ribosome type in PGCs. Therefore, maternal ribosomes may comprise a distinct set of ribosomes that drive translation in the germline - a finding that may have broad implications for understanding germline determination and development in zebrafish.

Heterogeneous rRNA sequences have been described in various organisms, including fungi, invertebrates (Dimarco et al. 2012), *Xenopus*, and humans (Pasolini et al. 2006; Vierna et al. 2013). However, whether ribosome heterogeneity translates into specialized ribosomal functions is still a matter of scientific debate. Apart from *Plasmodium* species (Gunderson et al. 1987; Li et al. 1997; Waters et al. 1997) and the 5S rRNA in *Xenopus* (Peterson et al. 1980), zebrafish embryos are the only known system that contains two distinct ribosomes composed of different sets of rRNAs. Differential expression of rRNA variants between tissues may indicate the ability of rRNA variant ribosomes to have distinct functions. Therefore, zebrafish embryogenesis, especially germline versus soma studies, provides a powerful system to study ribosome-mRNA interactions in the context of the functional consequences of ribosome heterogeneity. However, the cell-to-cell variability in their abundance requires future techniques to address these questions at the single cell level.

Importantly, we observed that at 1 dpf, when maternal and somatic ribosomes are present at similar levels, both types of ribosomes associate with polysomes and are thus likely involved in translation. Consistent with the role of maternal ribosomes in supporting early development, translation by maternally deposited ribosomes is essential and increases during embryogenesis before the onset of ribosome biogenesis (Bazzini et al. 2016; Hensey and Gautier 1997; Leesch and Lorenzo-Orts et al. 2023). Notably, in nematode worms lacking zygotic production of ribosomes (Cenik et al., 2016) and in zebrafish mutants with impaired ribosome biogenesis (Bielczyk-Maczyńska et al. 2015; Noack Watt et al. 2016), maternal ribosomes have been shown to be sufficient to sustain embryonic development.

If both maternal and somatic ribosomes are capable of translation, an obvious question is why a dual ribosomal system has evolved in zebrafish. One hypothesis is that maternal ribosomes may favor the translation of certain transcripts. Fully addressing this question would require swapping the maternal and somatic rRNA locus, a challenging experiment in zebrafish. Given that the major sequence differences between maternal and somatic rRNAs are in expansion segments, transcript specificity (if any) may be caused by solvent-exposed surfaces of maternal and somatic ribosomes. Our finding that 80S “hybrid” monosomes composed of 40S and 60S subunits of different origins are readily formed during zebrafish embryogenesis argues against maternal and somatic ribosomes acting as distinct entities.

As previously mentioned, though many sequence differences exist between the two, compensatory mutations of Watson-Crick base-pairs may preserve rRNA secondary structure in maternal and somatic ribosomes. For instance, both strands of helix 12 (H12) of the 28S rRNA differ in maternal and somatic ribosomes, with the somatic variant containing a U (U68) and the maternal sequence containing a C (C68). In the opposite strand, the maternal variant encodes a compensatory G73, whereas the somatic variant contains A73, thereby maintaining the Watson-Crick interaction (U-A in the somatic rRNA, C-G in the maternal rRNA). The sarcin-rich domain (SRD, or sarcin-rich loop, SRL) (Anger et al. 2013; Khatter et al. 2015) is a functionally important site of the ribosome, and contains two nucleotide differences (U_som_3716/C_mat_3864 and A_som_3738/G_mat_3886) located at the helix that forms the base of the SRL stem-loop. It is unclear if this represents a molecular difference among either ribosomes’ capacity to anchor elongation factors during tRNA translocation (Shi et al. 2012; Szewczak et al. 1993). Other functionally important regions like the GTPase association center (GAC), which acts as a binding platform for elongation factors and ribosome rescue factors (Uchiumi and Kominami 1994), were not resolved in our maternal ribosome cryo-EM map in the region containing three nucleotide differences between maternal and somatic 28S RNAs (A_som_1558/C_mat_1573, U_som_1560/C_mat_1575 and U_som_1565/C_mat_1580). Therefore, future studies will be required to directly compare the structures of maternal and somatic ribosomes, though our work already reveals that the vast majority of nucleotide differences is located at the surface on the ribosome and thus unlikely to alter for example ribosomal catalysis.

An alternative and non-exclusive hypothesis is that the maternal rDNA locus may be involved in PGC fate and sex determination in zebrafish. Laboratory strains of zebrafish lack sex chromosomes and, unlike mammals, rely on the transfer of specific maternal factors for germ cell development. Several reports suggest a role for the maternal rDNA locus in sex determination, with specific changes (such as demethylation and amplification on extrachromosomal circles) occurring during oocyte expansion and being associated with feminization (Breit et al. 2020; Dranow et al. 2013; Ortega-Recalde et al. 2019; Tao et al. 2020).

Another possibility is that the distinct set of rRNAs affects ribosome stability, as it has been suggested that maternal ribosomes are degraded more rapidly than their somatic counterparts (Locati et al. 2017b). A recent study proposed that ribosome ubiquitination plays a role in targeting and degrading maternal ribosomes during zebrafish embryogenesis (Ugajin et al., 2023). Additionally, one of the expansion segments, ES4L, is located on the ribosome surface and is formed by the 3’ end of the 5.8S rRNA and the 5’ end of the 28S rRNA. This 3’ half of the 5.8S rRNA has previously been suggested to play a role in ribosome degradation (Locati et al., 2018), and therefore this region could potentially serve as a target for degradation of maternal ribosomes.

Taken together, our study describes a distinct set of ribosomes present in the zebrafish germline. The presence of two distinct ribosomes during zebrafish embryogenesis provides an experimentally accessible system for future studies of ribosome specialization and defines a potentially novel mechanism of translational regulation in the zebrafish germline.

## Materials and Methods

### Zebrafish lines and husbandry

Zebrafish (*D. rerio*) experiments in the Pauli lab were conducted according to Austrian and European guidelines for animal research and approved by local Austrian authorities (animal protocols for work with zebrafish: GZ 342445/2016/12 and MA 58-221180-2021-16), and zebrafish lines in the Calo lab were housed in AAALAC-approved facilities and maintained according to protocols approved by the Massachusetts Institute of Technology Committee on Animal Care (CAC). Zebrafish transgenic lines containing maternally or somatically tagged ribosomes were generated as part of this study and are described below. In vivo samples were allocated randomly to the experiment and treated equally.

TLAB fish, generated by crossing zebrafish AB with the natural variant TL (Tupfel Longfin), served as wild-type zebrafish for experiments in the Pauli lab (**Figures 1B, 2, 3, and Supplementary Figures S2, S5 and S6**). All experiments in the Calo Lab were performed in the AB/Tübingen (TAB5/14) zebrafish genetic background (**Figures 4, 5, 6, and Supplementary Figures S1B, S4, S7, S8 and S9**). Fish were raised according to standard protocols (28 °C; 14:10 h light:dark cycle).

### Zebrafish embryo and tissue collection

Zebrafish embryos were collected at the indicated times post-fertilization, and staged according to (Kimmel et al. 1995).

Zebrafish AB9 fibroblasts, originally derived from an adult caudal fin (ATCC CRL-2298), were cultured on cell culture-treated 10 cm plates (Genesee Scientific, 25-500), collected using trypsin (Gibco, 25200072) and standard cell culture procedures. AB9 cells were grown at 28 °C with 5% CO_2_ in Dulbecco’s Modified Eagle Medium (DMEM, Genesee Scientific, 25–500) supplemented with 10% fetal bovine serum (FBS, Gemini Bio-products) and 1% Penicillin/Streptomycin (Gibco, 10378–016).

Zebrafish ZMEL melanoma cells, originally derived from Tp53^M214K/M214K^ tumors expressing human oncogenic BRAF^V600E^ were cultured and collected using the same protocols as AB9 cells. ZMEL cells were a kind gift from the laboratory of Dr. Leonard Zon (Boston Children’s Hospital).

Zebrafish ZF4 fibroblasts, originally derived from a 1 dpf embryo (ATCC CRL-2050), were cultured on cell culture-treated 10 cm plates (Genesee Scientific, 25-400), collected using trypsin (Gibco, 25200072) and standard cell culture procedures. ZF4 cells were grown at 28 °C with 5% CO_2_ in a 1:1 mixture of Dulbecco’s Modified Eagle Medium and Ham’s F-12 media (DMEM/F12, Genesee Scientific, 25-503) supplemented with 10% fetal bovine serum (FBS, Gemini Bio-products) and 1% Penicillin/Streptomycin (Gibco, 10378–016).

For all cell lines, cells from a 50-75% confluent 10 cm plate were collected and either flash frozen or directly lysed in TRIzol.

Zebrafish tumors used in this study were derived from the genotypes Tg(mitfa: GNAQ^Q209L^); Tp53^M214K/M214K^ (referred to as GNAQ melanoma), Tg(BRAF^V600E^); Tp53^M214K/M214K^ (referred to as BRAF melanoma), and Tp53^M214K/M214K^ (referred to as MPNST, malignant peripheral nerve sheath tumor) and excised using standard protocols approved by the Committee on Animal Care at MIT.

Adult zebrafish were monitored and euthanized before tumors (3-4 mm in diameter) were excised with a sterile scalpel and placed into a 35 mm dish. Tumors were incubated with 2 mL of Dissection Media [Dulbecco’s Modified Eagle Medium (DMEM, Genesee Scientific, 25–500), 1% Penicillin/Streptomycin (Gibco, 10378–016), and 75 μg/mL of Liberase (Roche, 5401020001)], manually disaggregated with a sterile razor blade, and let sit at room temperature until a cellular slurry formed. Tumors were further broken down by trituration with an additional 5 mL of Wash Solution [1x PBS (Genesee Scientific, 25– 508), 1% Penicillin/Streptomycin (Gibco, 10378–016), and 10% fetal bovine serum (FBS, Gemini Bio-products)]. Cells were passed through a 40 μm strainer (Falcon, 352340), pelleted for 2 min at 500 xg, washed twice with 1x PBS, pelleted, and lysed in TRIzol.

### Generation of transgenic fish lines

Plasmid construction was based on the Tol2/Gateway zebrafish kit (Kwan et al. 2007). The p5e-ubi entry clone was a generous gift from the laboratory of Dr. Leonard Zon (Boston Children’s Hospital). The p5e-sycp plasmid was cloned by amplifying the sycp promoter from genomic DNA before using Gateway BP Clonase II to insert the gel-selected product into pDONRP4-P1R (#219) using previously described primers (Gautier et al. 2013). The pMe-3xFLAG-eGFP-Rpl10a plasmid was cloned by stitching together PCR products containing 3xFLAG, eGFP, and Rpl10a coding sequence (amplified from 1 dpf cDNA) via Gibson Assembly (Gibson et al. 2009), then amplified using gateway primers before using Gateway BP Clonase II (ThermoFisher Scientific, 11789020) to clone the insert product. The pMe-3xFLAG-eGFP-Rpl10a-430-3’UTR plasmid was similarly cloned. The p3e-polyA entry clone (#302) and pDestTol2 (#393) were used from the Tol2Kit in conjunction with the above plasmids and LR Clonase II (ThermoFisher Scientific, 11791020) to generate ubi:eGFP-3xFLAG-Rpl10a:polyA and sycp:eGFP-3xFLAG-Rpl10a-430-3’UTR:polyA and are referred to as Tg(Som-RiboFLAG) and Tg(Mat-RiboFLAG) respectively.

To generate mosaic transgenic fish lines, a 1 nL mixture of plasmid DNA (40 ng/μL) and capped-mRNA encoding Tol2 Transposase (20 ng/μL) was loaded into needles pulled from glass capillary tubing (Warner, 64–0766) and injected into one-cell-stage embryos using a pico-liter injector (Warner Instruments, PLI90A). Mosaic founders were crossed with wild-type animals to generate a stable line.

F3 and F4 embryos were used for imaging and experiments. Brightfield images and eGFP images were acquired on a Leica M205 FCA fluorescence stereo microscope. Images were taken with the same settings for all embryos.

### FACS isolation of primordial germ cells (PGCs)

One-cell-stage zebrafish embryos were injected with 1-2 nL of capped *eGFP-nanos3’UTR* mRNA using a pico-liter injector (Warner Instruments, PLI90A) and needles pulled from glass capillary tubing (Warner, 64–0766). At 24 hpf, embryos were sorted for eGFP expression, dechorionated using 1 mg/mL pronase (Pronase from *Streptomyces griseus*, Cat# 000000010165921001, Sigma-Aldrich), and washed with 1x PBS (Genesee Scientific, 25–508).

80 eGFP-positive embryos were incubated in a 5 mL solution of TrypLE Express (ThermoFisher Scientific, 12605036) and 0.003% Tricaine (ethyl-3-aminobenzoate-methanesulfonate) for 60 min at 31 °C in a Thermomixer (Eppendorf, Thermomixer R) set to 100 rpm, followed by trituration using a glass Pasteur pipette until tissues were digested into single cells. Cells were passed through a 40 μm strainer (Falcon, 352340), pelleted for 5 min at 500 xg at room temperature, and washed with 1x PBS. Cell pellets were resuspended in 1x PBS supplemented with 5% fetal bovine serum (FBS, Gemini Bio-products). Using a BD FACSAria cell sorter, approximately 1,200 eGFP-expressing cells (primordial germ cells) and 1,200 non-eGFP-expressing cells (somatic cells) were sorted directly into TRIzol.

### Additions to polysome gradient protocols

To obtain a better resolution of the low molecular weight fractions, 24 hour post-fertilization embryos were batch deyolked according to the previously published protocol (Link et al. 2006). While 200 to 250 embryos were used per sample for non-deyolked samples, 500 embryos were used per gradient for deyolked embryos. The deyolking buffer was supplemented with 1x protease inhibitors (EDTA-free, Roche) and 0.1 μg/mL cycloheximide. To avoid DNA contamination in the higher molecular weight factions, obtained larval lysates were treated with 1:125 Turbo DNase (QIAGEN, 79254) according to the manufactures protocol and incubated for 10 min at 28 °C.

To isolate RNA after fractionation, 400 μL from each collected fraction were transferred into a Safe-lock Eppendorf tube, 1000 μL TRIzol (ThermoFisher Scientific, 15596-018) reagent were added and samples were vortexed for 10 s immediately. The following steps were performed according to the standard TRIzol based RNA isolation protocol.

### Western blotting of polysome fractions

400 μL of each fraction was first mixed with 700 μL water to dilute sucrose. Sodium deoxycholate and TCA were added to a final concentration of 0.08% and 20% respectively. Samples were vortexed and incubated overnight at −20 °C and centrifuged at 20,000 xg at 4 °C for 30 min. Protein pellets were washed twice with 2 volumes of cold (−20 °C) 100% acetone, then pelleted again. After acetone was removed, protein pellets were air dried, resuspended in 100 μL Laemmli buffer, sonicated for 5 min (30 s on, 30s off), and boiled for 5 min, before running a portion on a 4-20% Tris-glycine polyacrylamide gel (Invitrogen, XV04205). Transfer to a PVDF membrane previously activated with methanol, was run in cold Tris-glycine transfer buffer (1x Tris-glycine, 20% methanol) at 80 V for 80 min. After transfer, membranes were blocked in 5% milk in PBST (1x PBS with 0.1% Tween-20) for 1 h and blotted with primary antibodies (1: 2000 anti-FLAG, Millipore Sigma, F3165, and 1:500 anti-Rpl7, ThermoFisher Scientific, A300-741A) overnight at 4 °C. Membranes were washed with PBST and blotted with secondary antibodies (1:10,000 anti-mouse, Invitrogen, 32230, or 1:10,000 anti-rabbit, Invitrogen, 32260) for 1 h at room temperature. All blots were sequentially probed SuperSignal West Femto (ThermoFisher Scientific, 34096) was used to develop the blot and a Bioanalytical Imaging System model c500 (Azure Biosciences) was used for imaging.

### Isolation of zebrafish embryos for total RNA time course

Embryos were snap-frozen in liquid nitrogen and stored at −80 °C. To maintain a uniform genetic background, all embryos for the time course experiment were collected from the same fish stock.

### Mapping of somatic rRNA differences onto the maternal rRNA structure

Nucleotide sequence alignments between maternal and somatic zebrafish rRNAs were generated in Clustal Omega (https://www.ebi.ac.uk/jdispatcher/msa/clustalo) (Sievers et al. 2011) for each of the rRNA classes (28S, 18S, 5.8S and 5S), and pairwise nucleotide sequence differences were identified. Nucleotide positions that differed in the somatic compared to the maternal version of the rRNA (see **Supplementary Table S2)** were highlighted via ChimeraX-1.6.1 (Goddard et al. 2018; Pettersen et al. 2021; Meng et al. 2023) on the rRNA structural model of the maternal 6 hpf ribosome (PDB: 7OYB, Leesch and Lorenzo-Orts et al., 2023). Moreover, 2D rRNA maps were downloaded from R2DT (Sweeney et al., 2021; https://rnacentral.org/r2dt) (somatic 18S, 5.8S, 5S rRNA 2D model: Danio rerio; 28S rRNA 2D model: Homo sapiens, with expansion segments that differed most in zebrafish modeled via the Vienna RNAfold package (Lorenz et al., 2011)), and used as basis for mapping and classifying maternal vs somatic nucleotide differences. Human rRNA 2D maps highlighting nucleotide position-specific Shannon Entropy values based on the Eukarya phylogeny were downloaded from Ribovision2 (https://ribovision2.chemistry.gatech.edu/; Bernier et al., 2014), and adapted by highlighting the positions of maternal-vs-somatic nucleotide differences in zebrafish (**Supplementary Fig S6B and S6C**).

### RNA extraction from TRIzol

Homogenized lysate was mixed with 0.2 mL chloroform per mL of TRIzol, vortexed, and centrifuged for 15 minutes at 16,000 xg at 4 °C. The upper aqueous phase containing RNA was mixed with an equal volume of 100% ethanol, vortexed, processed with a Zymo column (RNA Clean and Concentrator, Zymo, R1014) including on-column DNase treatment, and eluted in 13 μL RNAse-free water.

### cDNA synthesis

cDNA was synthesized using SuperScript IV VILO Master Mix (ThermoFisher Scientific, 11766050). For each sample, an 8 μL reaction composed of 6.4 μL RNA, 0.8 μL 10x ezDNase Buffer, and 0.8 μL ezDNAse enzyme was incubated at 37 °C for 5 min before centrifuging and placing on ice. Then, 3 μL of SuperScript IV VILO Master Mix or No RT Control and 4 μL of RNase-free water was added to each reaction, followed by incubation at 25 °C for 10 min, 60 °C for 10 min, and 85 °C for 5 min.

### Detection of maternal and somatic rRNA variants by PCR

Primer sequences and annealing temperatures are listed in **Supplementary Table S3**. For certain PCRs, due to the high G/C content of rRNAs and therefore of the corresponding cDNA, standard PCR conditions were adapted to these more complex templates. RNA and cDNA were isolated as described above. The PCR reaction differed from the manufacturer’s protocol in the addition of 1.3x Q5® Hot Start High-Fidelity (from a 2x MasterMix, New England Biolabs M0494) and additionally 1 M betaine (Sigma-Aldrich, B0300). The PCR conditions were modified according to the standard protocol by extending of the elongation time up to 90s in each PCR cycle and a final elongation step of 15 min.

### Detection of maternal and somatic rRNA variants by RT-qPCR

The synthesized cDNA was diluted 5-1000 fold with RNase-free water, depending on the input material, and used in a 10 μL RT-qPCR reaction composed of 5 μL 2x KAPA SYBR FAST qPCR Master Mix (Roche, KK4600), 2.5 μL cDNA, and 0.25 μM of each primer. Reactions were arrayed into a 384-well reaction plate or a 96-well reaction and measured with a Light Cycler 480 II Real-time PCR Machine: 95 °C for 30 s, 35 cycles of 95 °C for 5 s, 60 °C for 15 s and 68 °C for 15 s. Three biological replicates, each from independent natural crosses, were measured in triplicate. No reverse-transcriptase samples were included as negative controls. Background amplification was determined using IPs from wild-type (uninjected) lysate. qPCR results with cycle numbers higher than the ‘background control’ were determined as ‘non-specific amplification’ and thus set to zero. Relative fold differences of specific targets were calculated using the 2ΔCt method. Statistical analyses were performed using Student’s unpaired t-test where p < 0.05 was considered significant.

### In vitro transcription of mRNA and biotin labeling

To generate the RNA for microinjections, 2 μg of a construct encoding eGFP-nanos3’UTR was digested using NotI-HF (New England BioLabs, R3189S) for 1 h. The linearized product was purified according to manufacturer specifications via gel selection using NucleoSpin (Macherey-Nagel, 740588.50). 500 ng of DNA was used in an overnight IVT reaction using mMESSAGE mMACHINE T7 ULTRA Transcription Kit (Invitrogen, AM1345). The product was treated with TURBO DNase for 30 min. RNA was poly-A-tailed using the provided tailing reaction and samples were run on agarose gels to confirm sizes. RNA was purified using Zymo RNA Clean and Concentrator columns (Zymo Research, R1016) and concentration was measured using a Nanodrop ND-1000 Spectrophotometer (ThermoFisher Scientific).

End labeling of RNA was achieved using previously described methods (Wang et al. 2007). Briefly, 20 pmol of in vitro transcribed mRNA (approximately 10 ug) was subjected to labeling with 1 nmol of pCp-Biotin (Jena Bioscience, NU-1706-BIO) in a 20 μL reaction comprising 5 μL DMSO, 2 μL 10 mM ATP, 1 μL T4 RNA Ligase 1 (NEB, M0437M), 1 μL SuperAseIn (Invitrogen AM2694), and 2 μL T4 RNA Ligase Buffer at 25 °C for 2 h. A no-labeling reaction was similarly incubated with the reaction components lacking pCp-Biotin. All mRNAs were purified using Zymo RNA columns and adjusted to 200 ng/μL.

### Isolation of zebrafish ribosome subunits by immunoprecipitation

Embryos were washed twice with 500 μL of ice-cold Subunit Buffer (20 mM Tris-HCl pH 7.5, 150 mM NaCl, 1 mM EDTA, and 1 mM DTT. As much buffer as possible was removed before samples were either immediately processed or flash frozen and stored at −80 °C.

400 μL of ice-cold Subunit Lysis Buffer (Subunit Buffer with 1% Triton X-100, 1x cOmplete-EDTA-free protease inhibitor (Sigma, 11873580001), and 20 mM EDTA) was added to either fresh or frozen tissue. Keeping tubes on ice, a pre-cooled 1 mL syringe attached to a 26-gauge needle was used to shear samples by slowly triturating material for 20-30 strokes until no visible tissue remained. Lysates were incubated on ice for 5 min, then cleared by centrifugation at 10,000 xg for 10 min at 4 °C.

Cleared S10 fractions were adjusted to 500 μL with Subunit Lysis Buffer and overlaid onto 2 mL of a 1 M sucrose cushion prepared in Subunit Buffer in a 15×51 mm 3 mL thickwall polycarbonate ultracentrifuge tube (Beckman Coulter, 349622) on ice. Ribosomal subunits were pelleted using a TLA100.3 rotor (Beckman Coulter, 349490) at ∼300,000 xg at 4 °C for 2 h. Supernatants were carefully poured off and pellets were resuspended by horizontal rotation on ice for at least 30 min in 235 μL of Subunit Buffer. Resuspended subunits were incubated with 2 μL of anti-FLAG antibody (Millipore Sigma, F3165-.2MG) for 1 h at 4 °C. Then, 14 μL of equilibrated Dynabeads Protein G (ThermoFisher Scientific, 10003D) were added and incubated at 4 °C for at least 4 h. Beads were subsequently collected using a magnetic rack, washed with Subunit Buffer three times, and resuspended in TRIzol-LS reagent (Invitrogen, 10296028).

### Isolation of zebrafish monosomes by immunoprecipitation

Embryos were washed twice with 500 μL of ice-cold Ribosome Buffer [20 mM Tris-HCl pH 7.5, 150 mM NaCl, 5 mM MgCl_2_, 1 mM DTT]. As much buffer as possible was removed before samples were either immediately processed or flash frozen and stored at −80 °C.

200 μL of ice-cold Ribosome Lysis Buffer [Ribosome Buffer containing 1% Triton X-100, 1x cOmplete-EDTA-free protease inhibitor (Sigma, 11873580001), and 100 μg/mL cycloheximide] was added to either fresh or frozen tissue. Keeping tubes on ice, a pre-cooled 1 mL syringe attached to a 26-gauge needle was used to shear samples by slowly triturating material for 20-30 strokes until no visible tissue remained. Lysates were incubated on ice for 5 min, cleared by centrifugation at 10,000 xg for 10 min at 4 °C, then treated with 1 unit per embryo of RNAse T1 (ThermoFisher, EN0541). After 90 min of rotating at room temperature, 2 units per embryo of SuperAseIn (Invitrogen, AM2694) were added to stop the reaction.

Cleared and digested S10 fractions were adjusted to 250 μL with Ribosome Lysis Buffer and overlaid onto a continuous 10-50% (w/v) sucrose gradient prepared in Ribosome Buffer containing 100 μg/mL cycloheximide. Gradients were generated using a BioComp Gradient Master device and 14×89 mm ultracentrifuge tubes (Seton Scientific, 7030). Gradients were centrifuged in a SW41 Ti rotor (Beckman Coulter, 331362) at 35,000 rpm (r_max_ of 210,000 xg) for 150 min at 4 °C followed by analysis and fractionation using a BioComp Gradient Station coupled to a Model Triax™ Flow Cell detector (FC-2). Approximately 1500 μL from the monosome fraction were manually collected.

The resulting monosomes were incubated with 2 μL of anti-FLAG antibody (Millipore Sigma, F3165-.2MG) for 1 h at 4 °C. Then, 14 μL of equilibrated Dynabeads Protein G (ThermoFisher Scientific, 10003D) were added and incubated at 4 °C for at least 4 h. Beads were subsequently collected using a magnetic rack, washed three times with Ribosome Buffer containing 100 μg/mL cycloheximide, and resuspended in TRIzol-LS reagent (Invitrogen, 10296028).

### RIP of biotinylated mRNAs

One-cell-stage zebrafish embryos were injected with 1-2 nL of either biotinylated or non-biotinylated *eGFP-nanos3’UTR* mRNA. At 24 hpf, embryos were dechorionated using pronase (Pronase from Streptomyces griseus, Cat# 10165921001, Sigma-Aldrich) and checked for eGFP expression.

Approximately 400 eGFP-positive embryos were washed twice with 500 μL of ice-cold Ribosome Buffer [20 mM Tris-HCl pH 7.5, 150 mM NaCl, 5 mM MgCl_2_, 1 mM DTT]. As much buffer as possible was removed before samples were immediately processed.

400 μL of ice-cold Ribosome Lysis Buffer [Ribosome Buffer containing 1% Triton X-100, 1x cOmplete-EDTA-free protease inhibitor (Sigma, 11873580001), and 100 μg/mL cycloheximide] was added to fresh tissue. Keeping tubes on ice, a pre-cooled 1 mL syringe attached to a 26-gauge needle was used to shear samples by slowly triturating material for 20-30 strokes until no visible tissue remained. Lysates were incubated on ice for 5 min, then cleared by centrifugation at 10,000 xg for 10 min at 4 °C.

Cleared S10 fractions were adjusted to 500 μL with Ribosome Lysis Buffer and overlaid onto 2 mL of a 1 M sucrose cushion prepared in Ribosome Buffer containing 100 μg/mL cycloheximide in a 15×51 mm 3 mL thickwall polycarbonate ultracentrifuge tube (Beckman Coulter, 349622) on ice. Ribosomes were pelleted using a TLA100.3 rotor (Beckman Coulter, 349490) at 80,000 rpm (r_max_ of 346,000 xg) for 2 h at 4 °C. Supernatants were carefully poured off and pellets were resuspended by horizontal rotation on ice for at least 30 min in 235 μL of Ribosome Buffer with 100 μg/mL cycloheximide and 5 μL SuperAseIn (Invitrogen AM2694).

Streptavidin magnetic beads (Invitrogen, 65001) were washed twice with Ribosome Buffer, then added to resuspended ribosomes, and incubated with rotation overnight at 4 °C. Beads were magnetically collected, washed three times with Ribosome Buffer containing 100 μg/mL cycloheximide and resuspended in TRIzol LS reagent (Invitrogen, 10296028).

### Data availability

Mass spectrometry proteomics data of isolated zebrafish ribosomes from 0, 1, 3 and 6 hpf has been published previously (Leesch and Lorenzo-Orts et al. 2023) and is accessible via the PRIDE partner repository with the dataset identifier PXD026866. Mass spectrometry proteomics data of isolated zebrafish ribosomes from 120 hpf (5 dpf) has been deposited to the ProteomeXchange Consortium via the PRIDE partner repository and will be made accessible upon publication. RNA-Seq data of total RNA (not ribosomal RNA depleted) was deposited to GEO and will be made accessible upon publication.

## Supplementary Figures

**Supplementary Figure S1:**
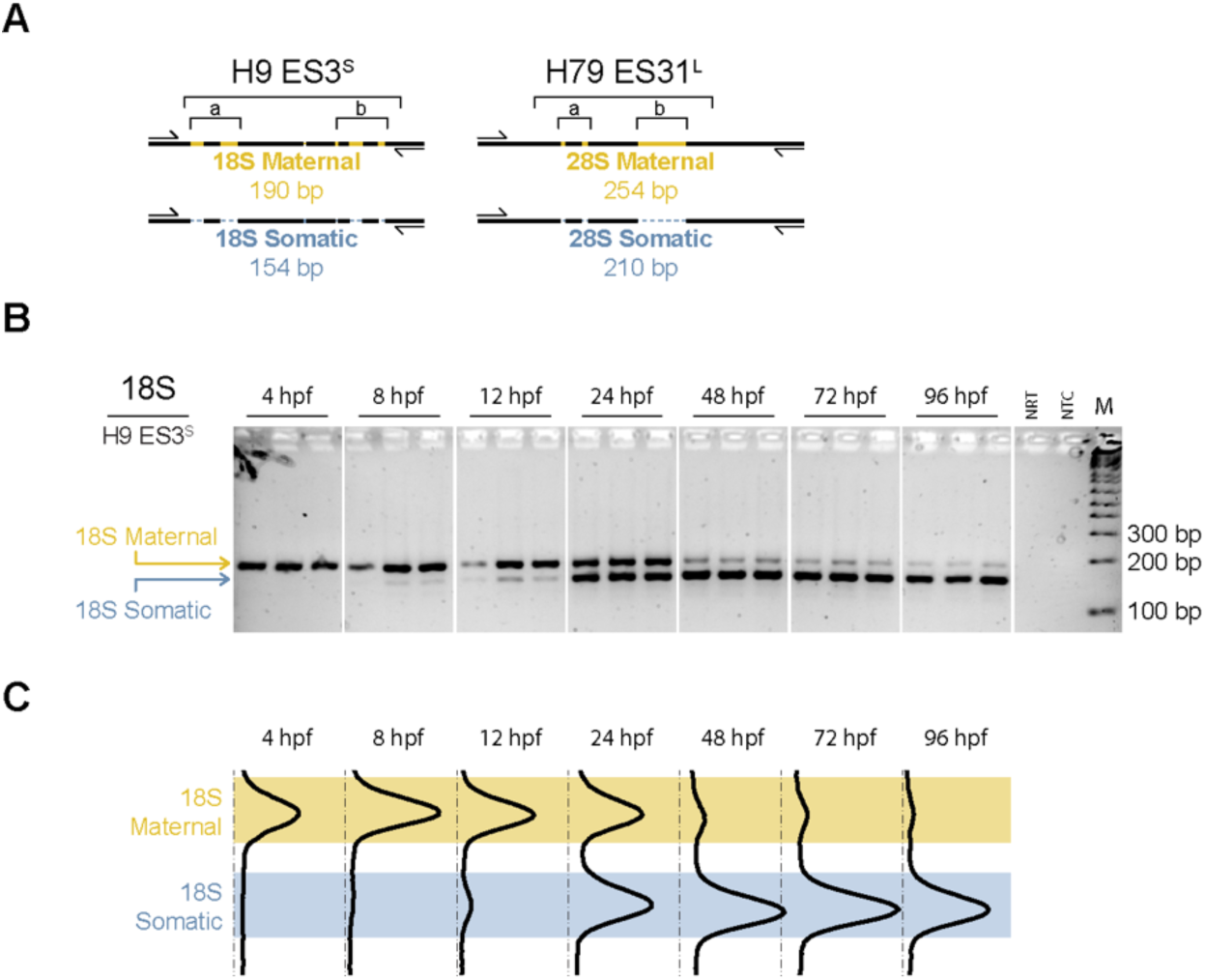
Detection of maternal and somatic rRNAs in embryonic development. **A)** Schematic representation of the rRNA expansion segments (ES) depicted in Fig. 1A which are targeted for a PCR-based fragment length polymorphism (FLP) assay. Either ES3S (helical regions *H9a* and *H9b*) in the 18S rRNA or ES31L (helical regions *H79a* and *H79b*) in the 28S rRNA are PCR amplified using a single primer pair (indicated by primer icons) detecting both maternal and somatic rRNA variants in the same reaction. Resulting PCR products differ in size due to the different lengths of ES regions in each rRNA. **Supplementary Table S3** contains the relevant PCR conditions and primer sequences. **B)** Gel electrophoretic analysis of PCR-based detection of maternal and somatic rRNA variants at the indicated developmental times. For each developmental time assayed, three individual embryos, each from an independent natural cross comprising one wildtype female and one wildtype male, were used for RNA extraction and cDNA synthesis. **C)** Averaged (n=3) line profiles of densitometry scans (ImageJ) from each lane of the gel shown in (B). The regions representing signal from maternal and somatic 18S rRNA PCR FLP are indicated and used for relative quantification in Fig. 1B. bp, base pair. NRT, no RT control. NTC, no template control. M, 100 bp marker.

**Supplementary Figure S2:**
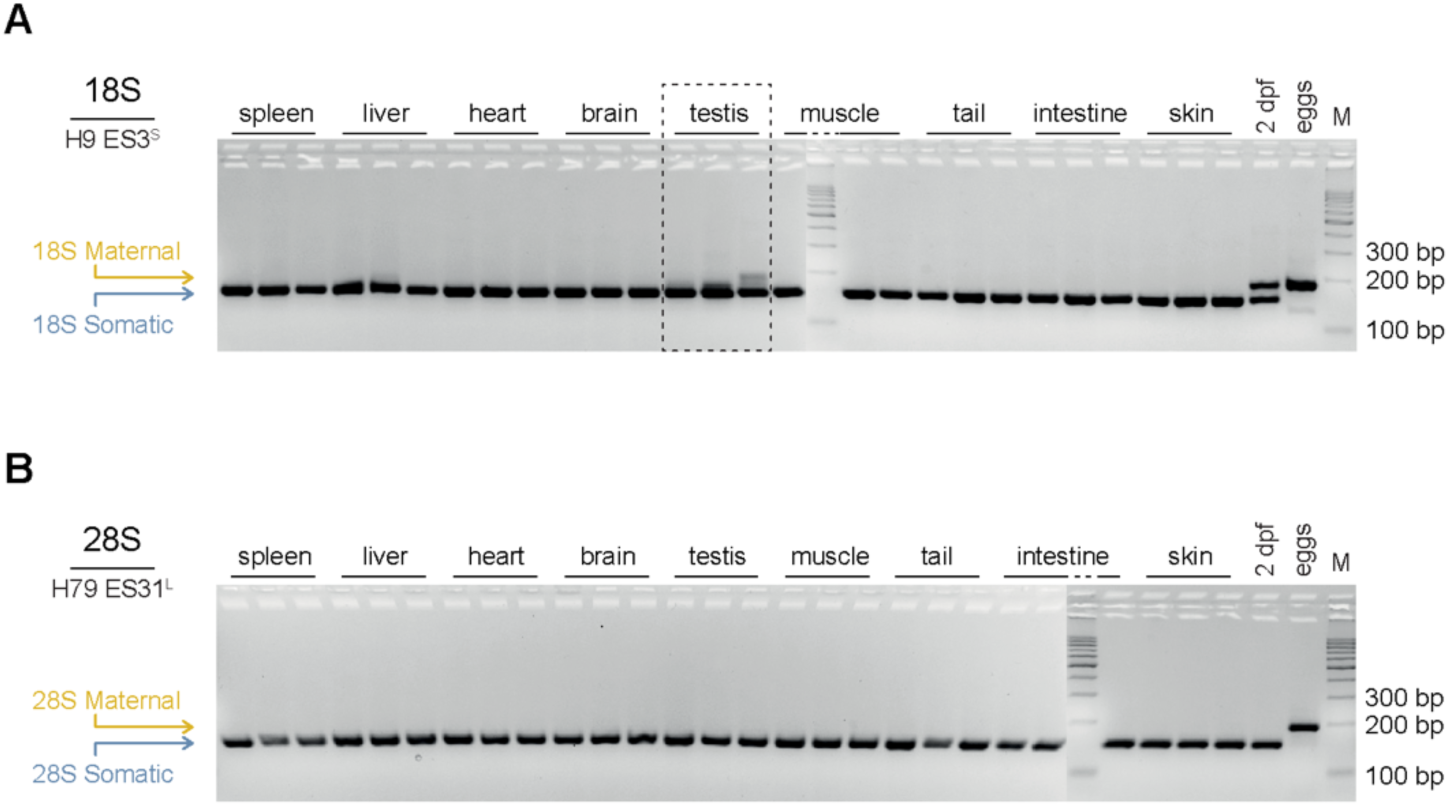
Analysis of rRNA variant expression in different tissues. Gel electrophoretic analysis of PCR-based detection of maternal and somatic **A)** 18S rRNA and **B)** 28S rRNA variants in different adult zebrafish tissues (n=3, each). Positive control reactions using 2 dpf and egg cDNAs are included for reference. See **Supplementary Fig. S1** and **Supplementary Table S3** for indicated lengths of PCR products from maternal and somatic rRNAs. H, helix. bp, base pair. M, 100 bp marker.

**Supplementary Figure S3:**
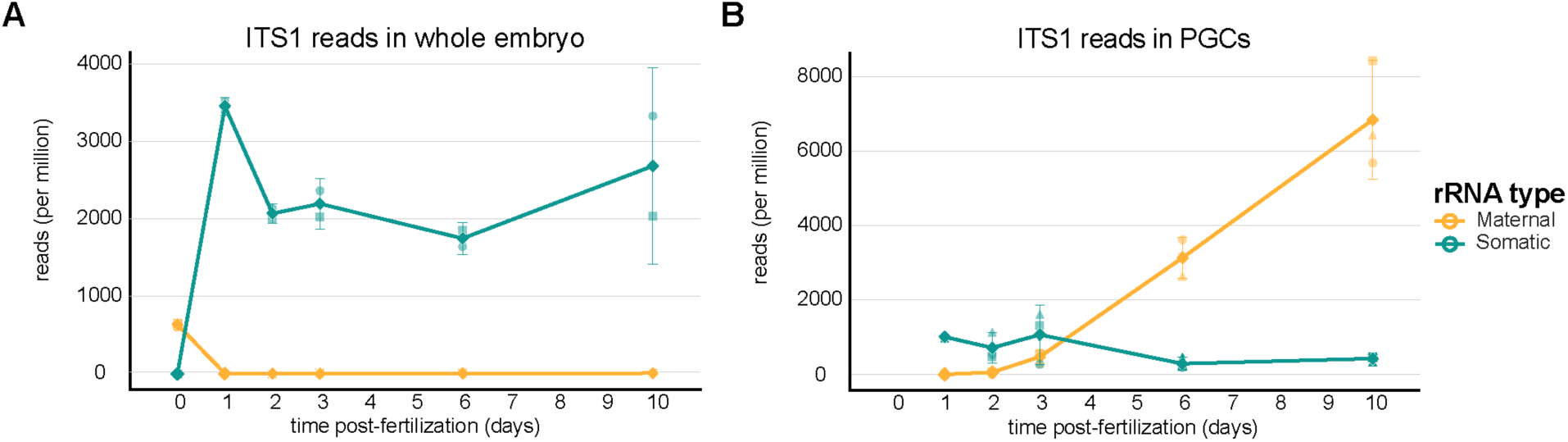
Transcription of rRNA variants in zebrafish and primordial germ cells (PGCs) from 1 through 10 days post fertilization. Expression of maternal (yellow) and somatic (blue) ITS1 sequences, indicative of de novo rRNA transcription, in **A)** whole embryos/larvae and in **B)** FACS-sorted PGCs. Public dataset PRJNA597223 (Redl et al. 2021) was obtained from the Sequence Read Archive at NCBI for analysis.

**Supplementary Figure S4:**
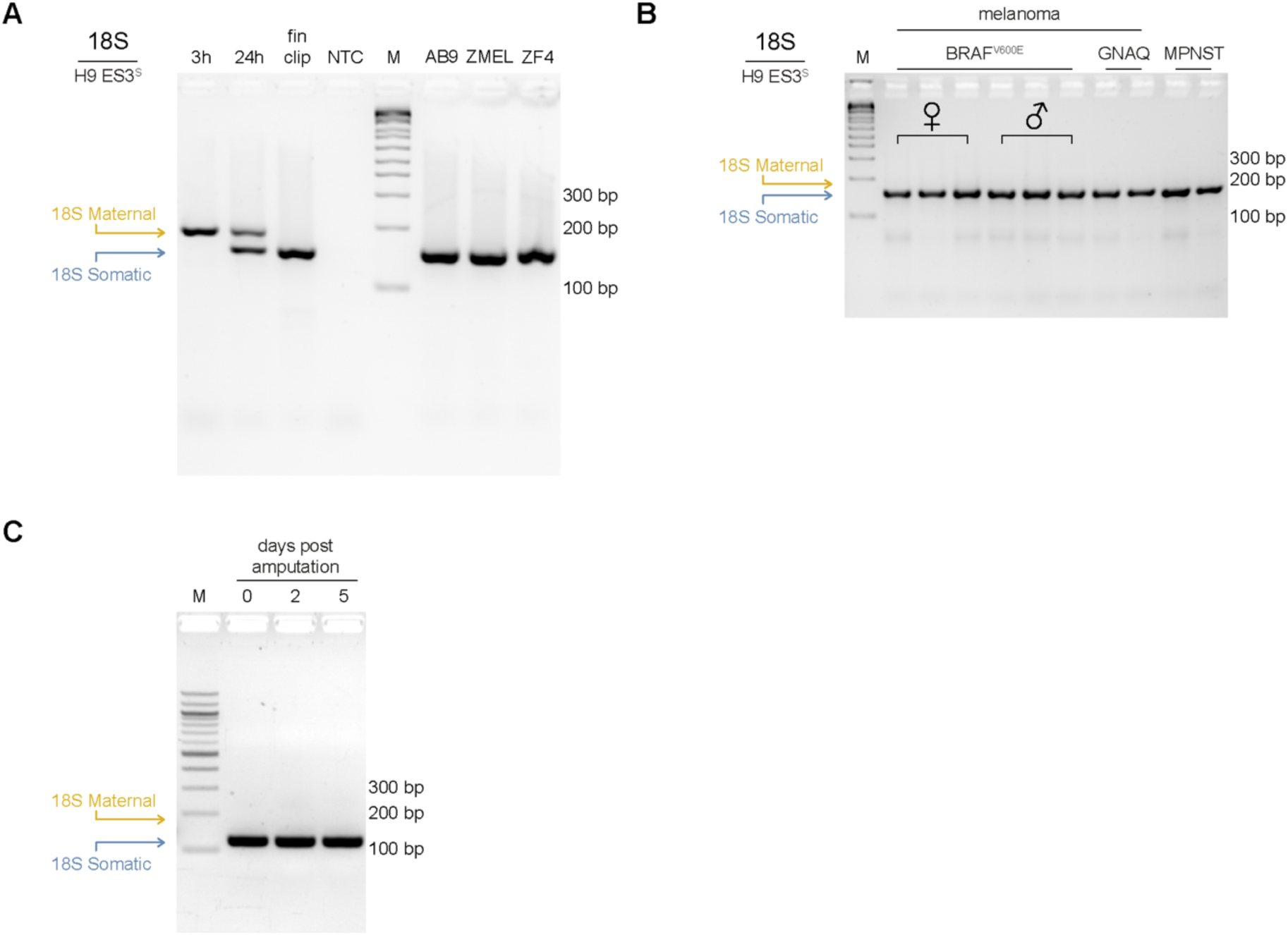
Detecting rRNA variants in cell lines, tumors, and regenerating fins. **A)** Gel electrophoretic analysis of PCR-based fragment length polymorphism (FLP) 18S rRNA variant detection from the indicated cell lines. Diagnostic PCR lengths for maternal and somatic 18S rRNAs are indicated. Positive control reactions using 3 hpf, 24 hpf, and adult fin clip cDNAs are included to compare maternal-only, maternal and somatic, or somatic-only band patterns, respectively. **B)** Gel electrophoretic analysis of 18S rRNA variant detection from the indicated excised tumors. See Materials and Methods for a complete description of each cell line and tumor. **C)** Gel electrophoretic analysis of the 18S rRNA FLP assay using cDNA made from regenerating fin blastema excised at the indicated times post initial amputation. NTC, no template control. AB9 (fibroblast cells), ZMEL (melanoma cells), ZF4 (fibroblast cells), BRAF (melanoma tumor), GNAQ (melanoma tumor), MPNST (malignant peripheral nerve sheath tumor).

**Supplementary Figure S5:**
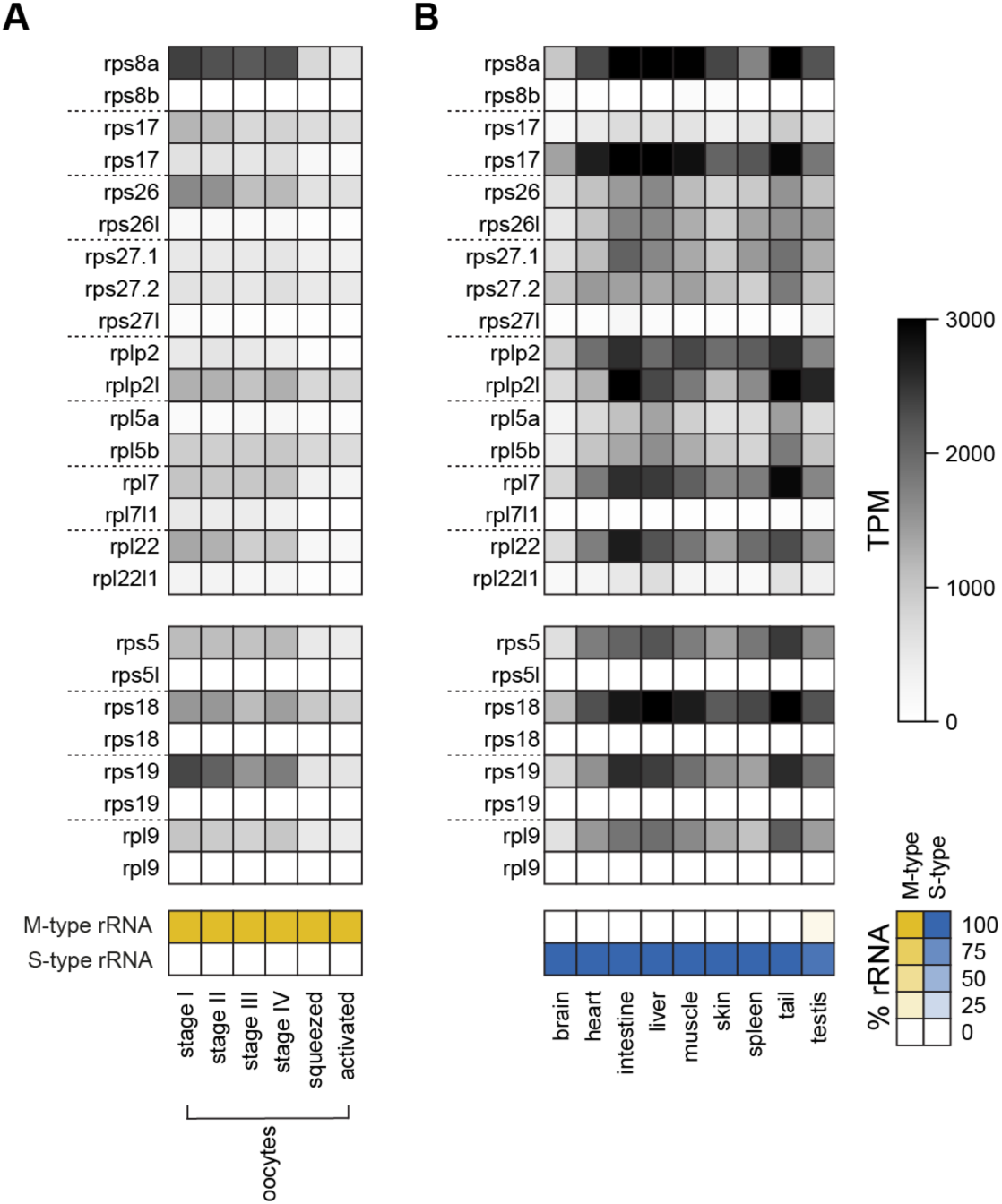
Expression of alternative ribosomal core protein mRNAs in zebrafish oogenesis and adult tissues. Heatmap of mRNA expression of paralogs and alternative isoforms encoding ribosomal protein variants assayed **A)** over progressive stages of oogenesis and **B)** across adult tissues and organs. The relative levels of maternal (yellow) or somatic (blue) rRNA variants at these developmental stages is indicated in the scheme below each heat map.

**Supplementary Figure S6:**
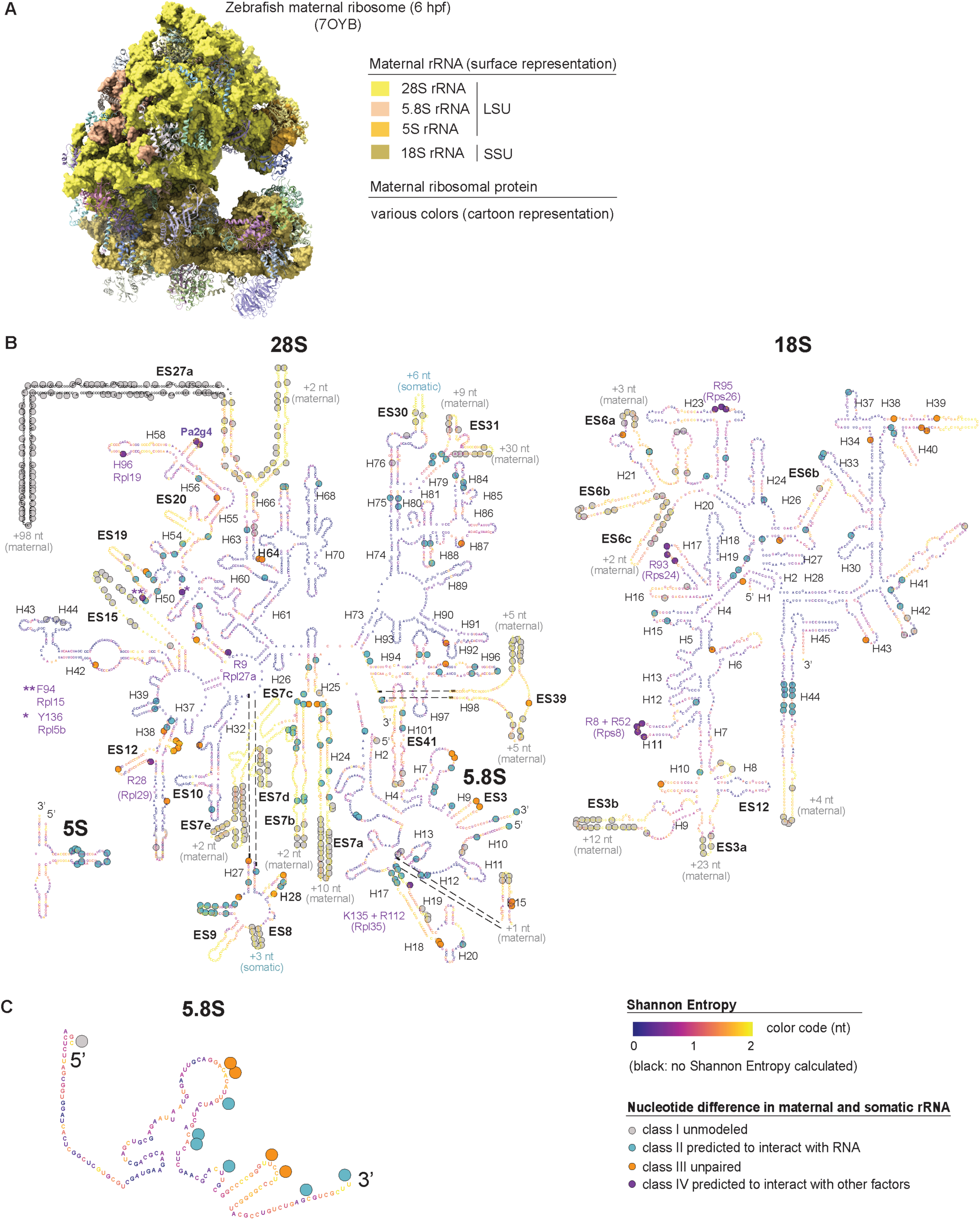
Sites that differ between maternal and somatic rRNAs in zebrafish are enriched in evolutionarily less conserved regions. **A)** Map of the maternal zebrafish ribosome isolated from 6 hpf embryos (PDB: 7OYB; Leesch and Lorenzo-Orts et al., 2023). Maternal rRNA sequences are shown in surface representation in shades of yellow; ribosomal proteins are shown in cartoon representation in various colors. **B)** Positions of sequence differences in zebrafish maternal and somatic rRNA variants mapped onto rRNA secondary structure predictions of human 28S, 18S, 5.8S and 5S rRNAs, color-coded for site-specific Shannon Entropy values, which were downloaded from Ribovision2 (https://ribovision2.chemistry.gatech.edu/; Bernier et al., 2014). A higher magnification view of the 5.8S rRNA is shown in C), highlighting the co-occurrence of evolutionarily variable nucleotides (high Shannon Entropy values) with sites that differ between zebrafish maternal and somatic rRNAs.

**Supplementary Figure S7:**
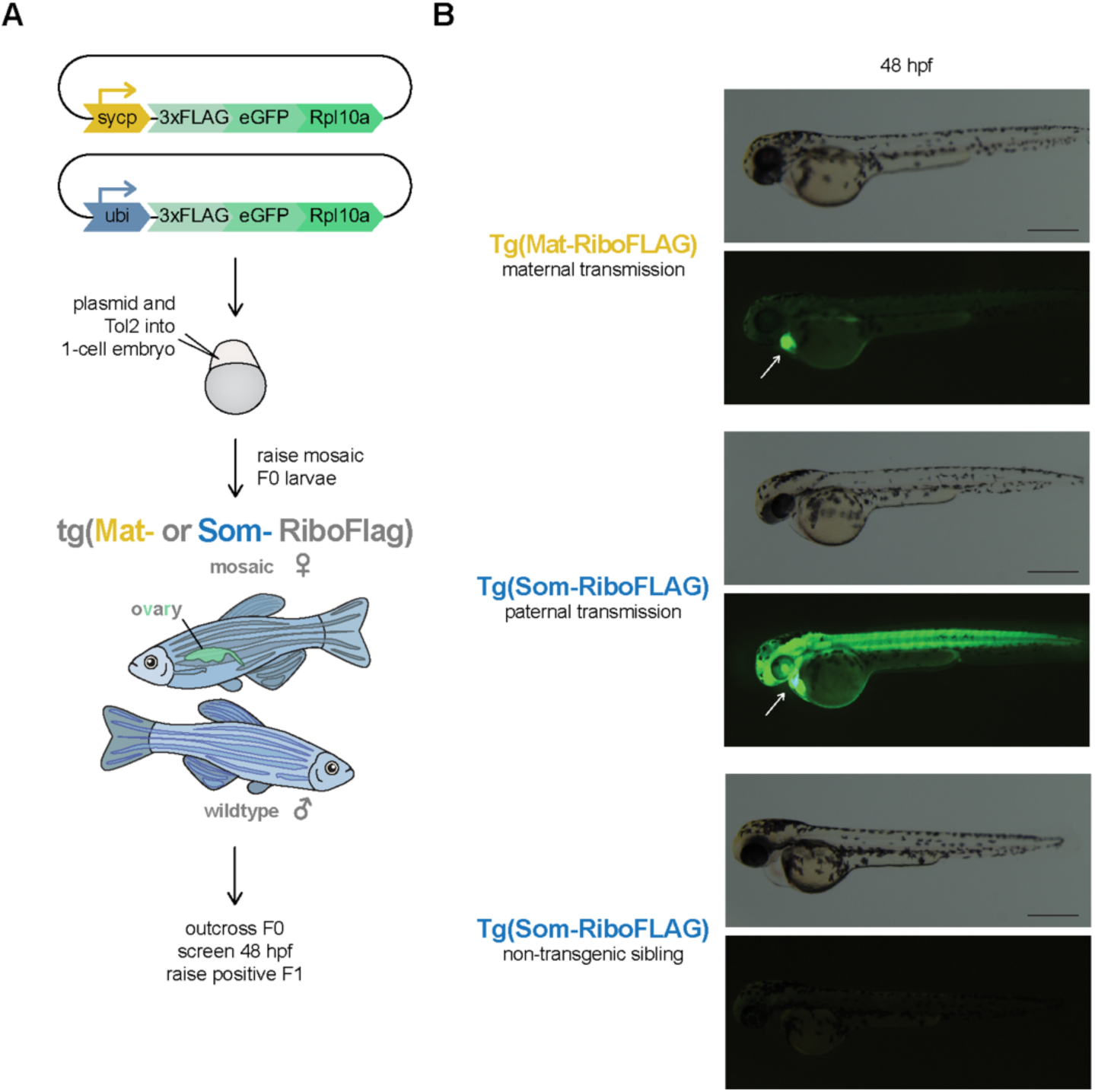
Generation of Tg(Mat-RiboFLAG) and Tg(Som-RiboFLAG) lines. **A)** Cartoon depicting the generation of two transgenic lines containing either FLAG-tagged maternal or FLAG-tagged somatic ribosomes (see Materials and Methods for details). Mosaic F0 were raised to adulthood and transmission of either transgene via the germline was screened for in **B)** 48 hpf embryos using a green heart marker (*cmlc2* promoter driving eGFP) also encoded on the plasmid (Kwan et al. 2007) as indicated by the white arrow. As expected, only half of the progeny derived from a Tg(Som-RiboFLAG) male and a wildtype female express the transgene. Scale bars indicate 500 μm. eGFP, enhanced green fluorescent protein. Tg, transgene.

**Supplementary Figure S8:**
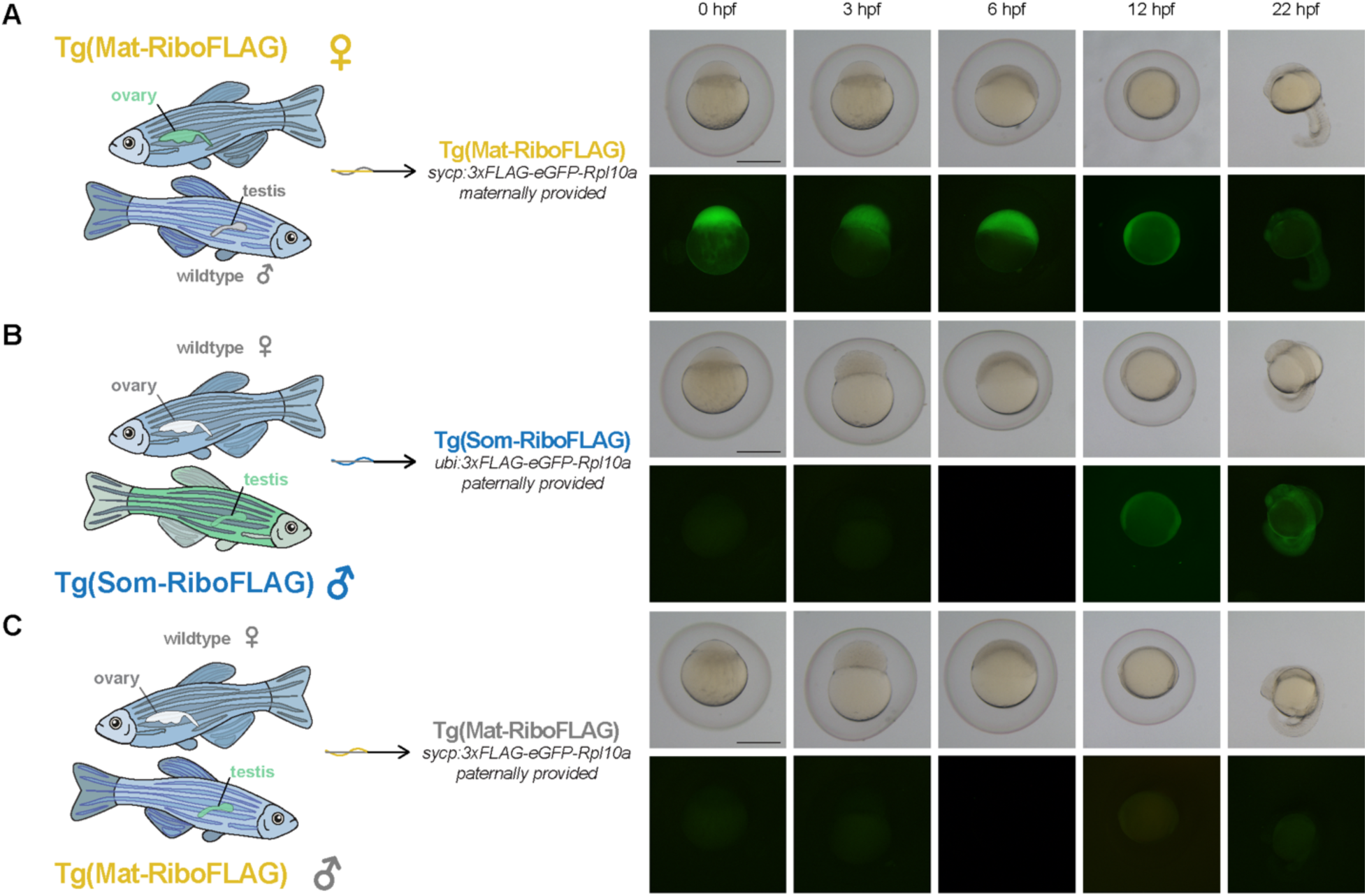
Transgenic expression 3xFLAG-eGFP-Rpl10a. Scheme summarizing the generation of embryos with FLAG-tagged ribosomes. Brightfield and fluorescent images of developing zebrafish embryos from either **A)** maternally-provided Tg(Mat-RiboFLAG), **B)** paternally-provided Tg(Som-RiboFLAG), or **C)** paternally-provided Tg(Mat-RiboFLAG) parents at the indicated hours post-fertilization. Embryos depicted in (A) contain maternally deposited 3xFLAG-eGFP-Rpl10a at 0 hpf while embryos depicted in (B) begin expression of 3xFLAG-eGFP-Rpl10a after 6 hpf. See **Supplementary Fig. S9** for expression at 48 hpf. As expected, progeny derived from paternally-provided Tg(Mat-RiboFLAG) show no transgene expression. Scale bars indicate 500 μm.

**Supplementary Figure S9:**
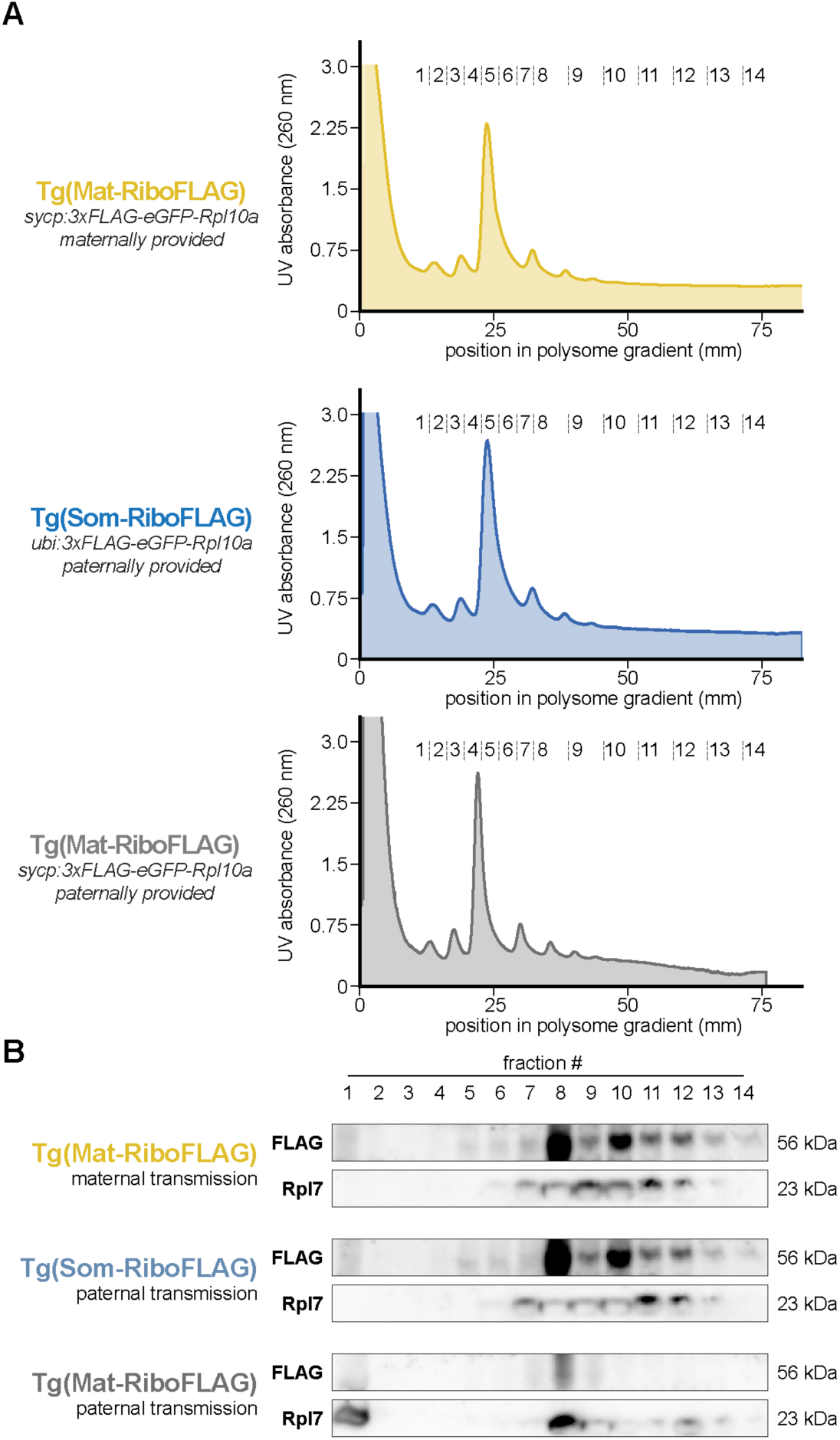
Incorporation of Tg(3xFLAG-Rpl10a) into translating ribosomes. **A)** Representative polysome profiles with continuous A260 reading from 24 hpf embryos. Position in the gradient and collected fractions are indicated. **B)** Western blots of polysome gradient fractions containing lysate from 24 hpf embryos with either maternally-provided Tg(Mat-RiboFLAG) or paternally-inherited Tg(Som-RiboFLAG) detecting the expression of transgenic FLAG-Rpl10a, and a negative control Western blot of polysome gradient fractions containing lysate from 24 hpf embryos with paternally-provided Tg(Mat-RiboFLAG) lacking the expression of transgenic FLAG-Rpl10a. kDa, kilodalton.

## Supplemental Tables

**Supplementary Table S1:**
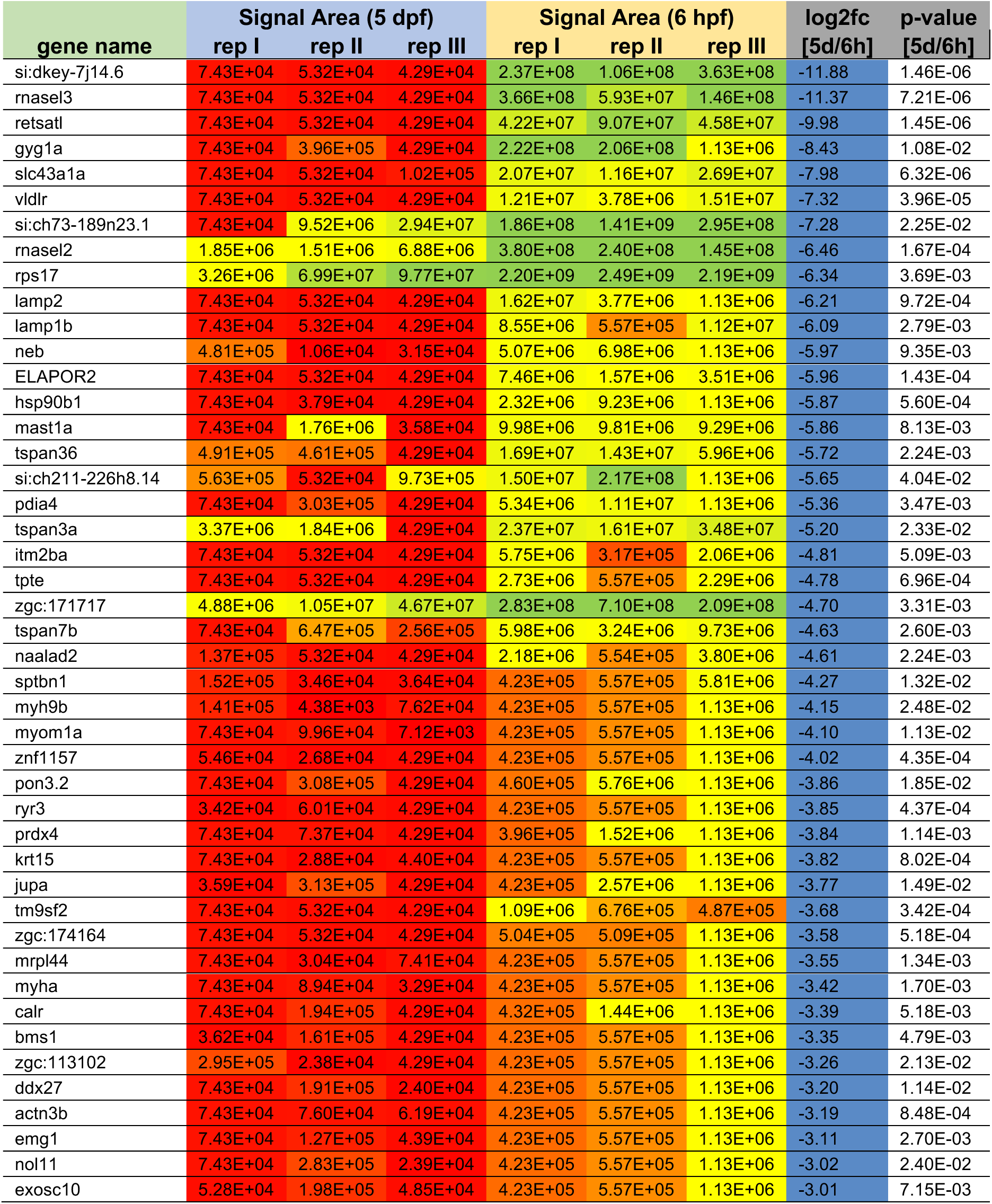
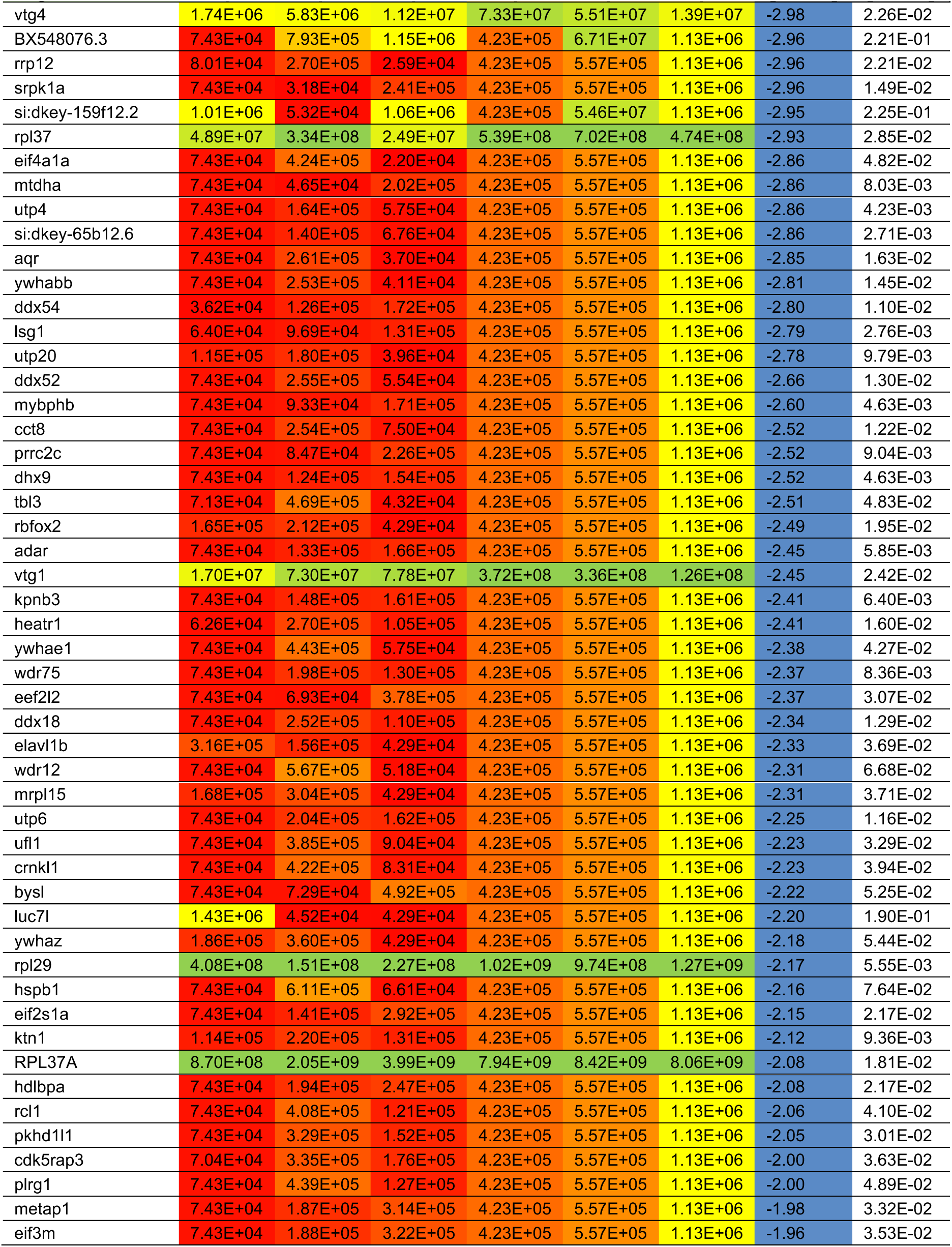
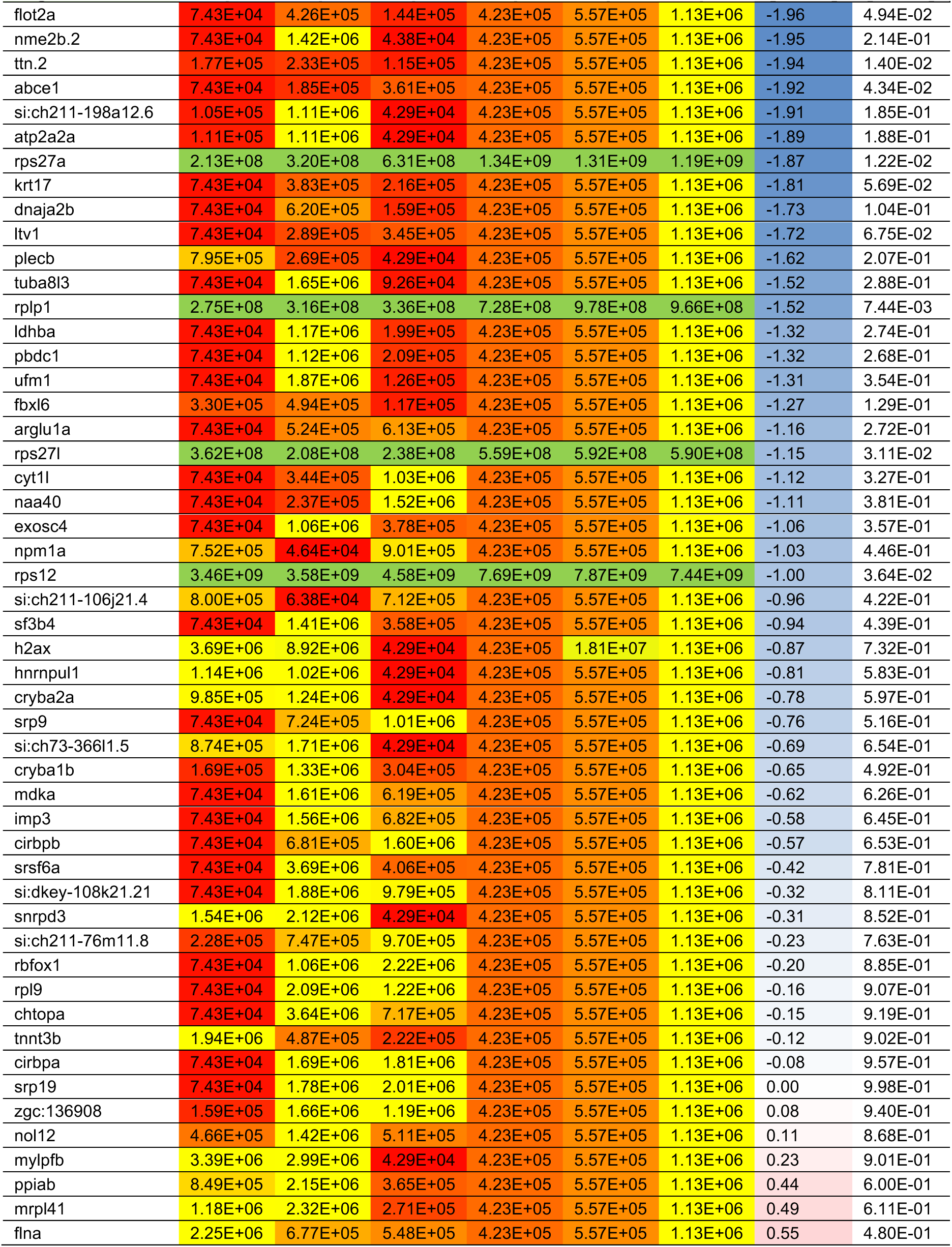
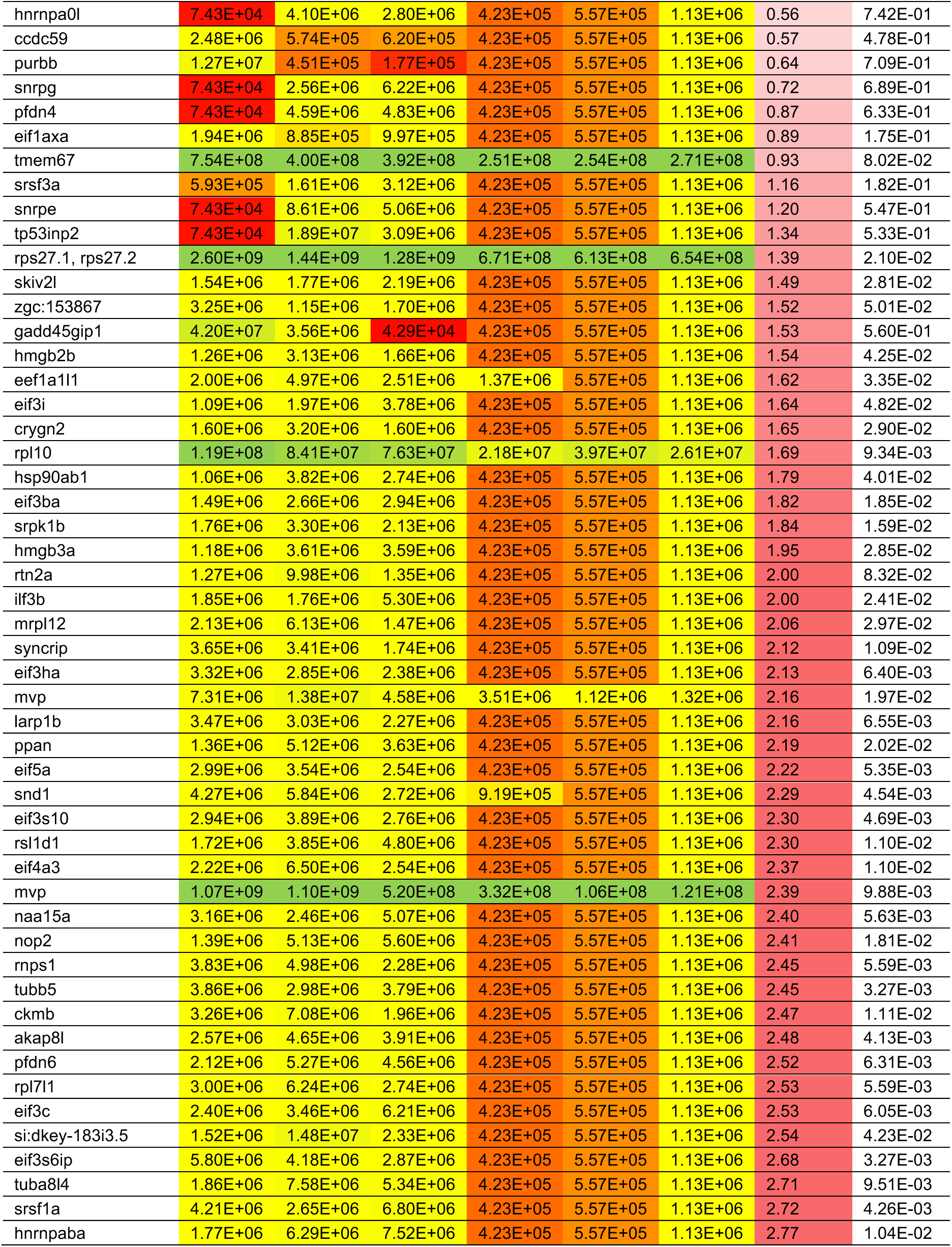
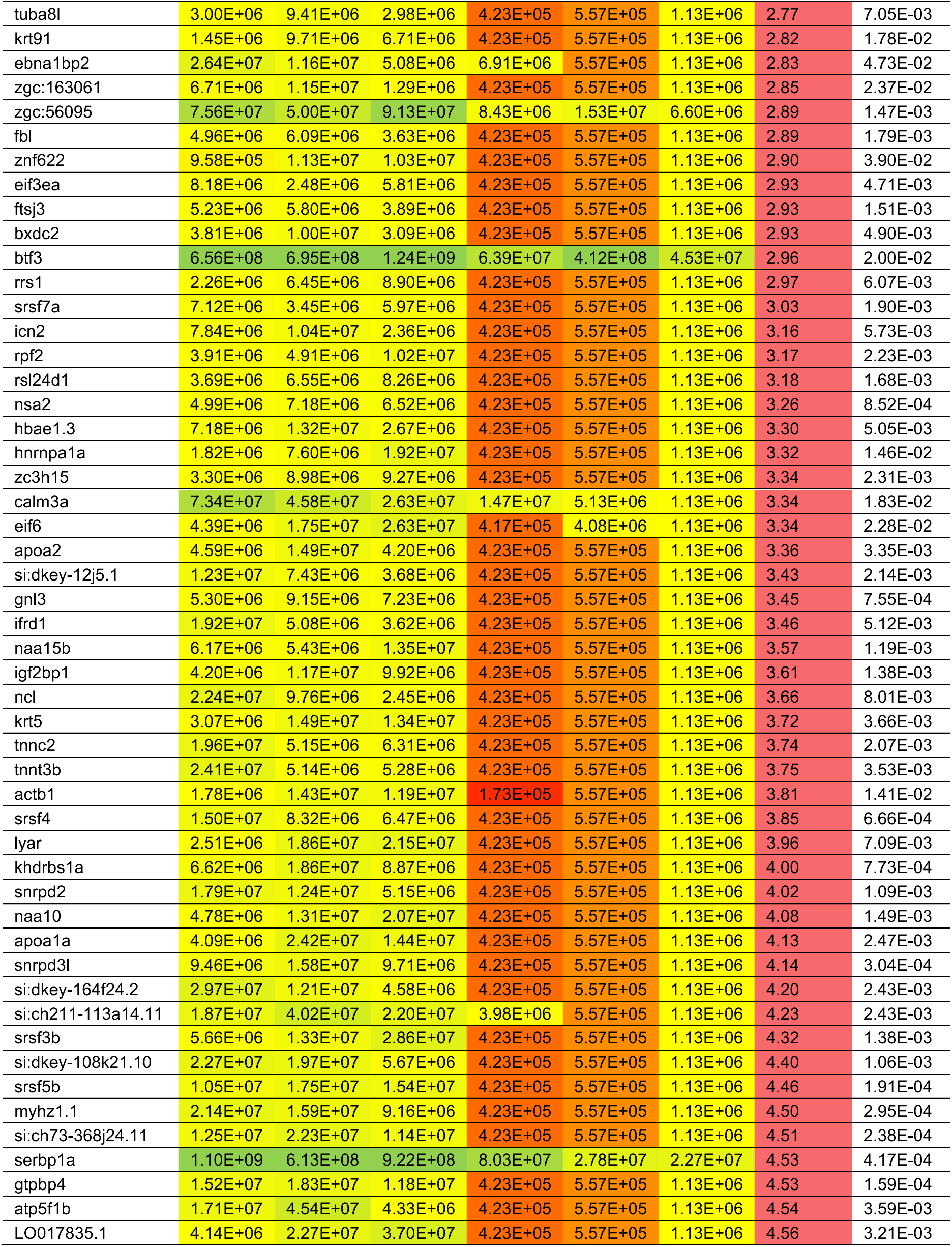
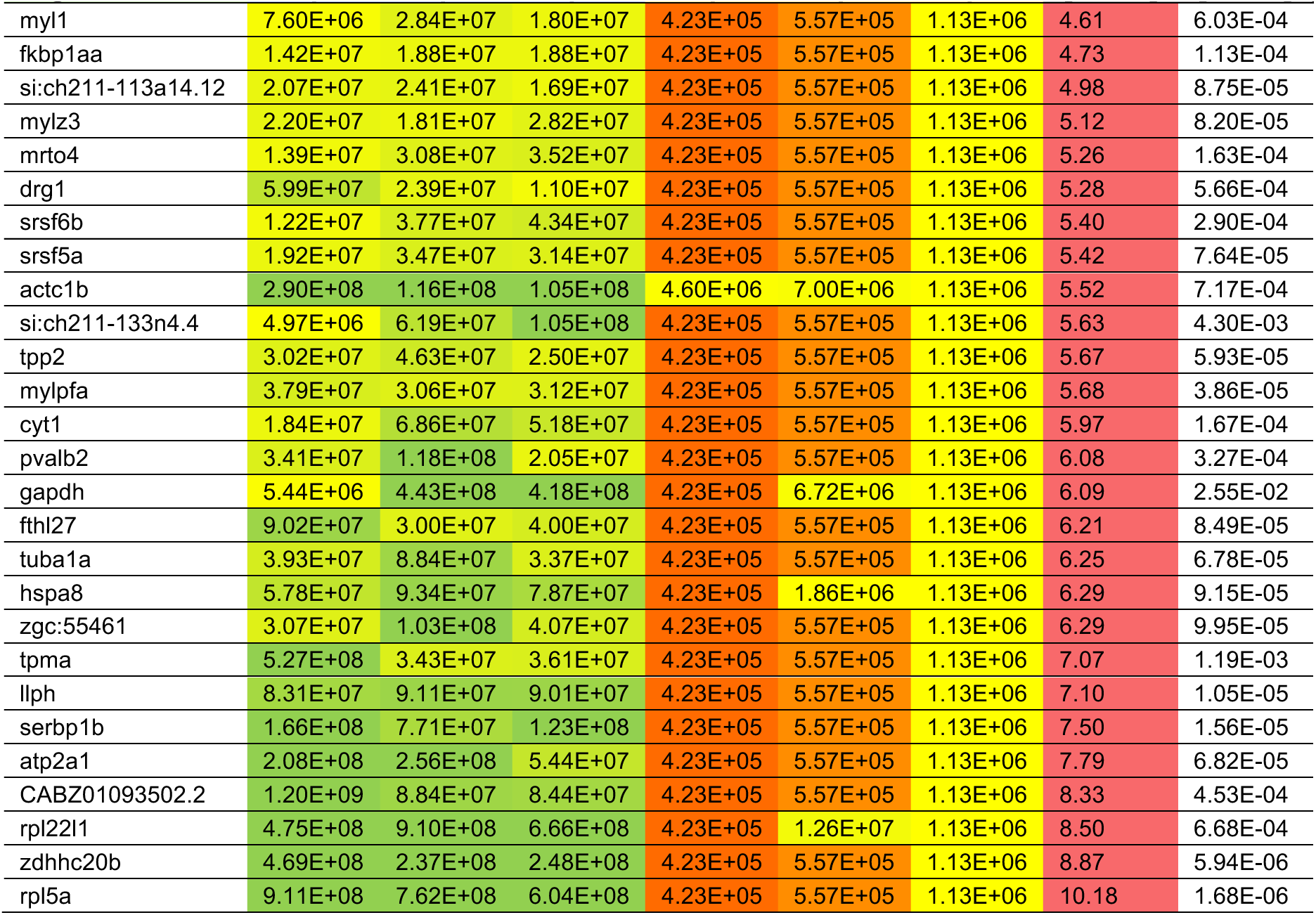
Proteins differentially associated with maternal and somatic zebrafish ribosomes. Proteins significantly enriched or depleted in ribosomes isolated from 6 hpf embryos versus 5 dpf larvae (permutation-based FDR <0.05).

**Supplementary Table S2:** Structural comparison of sequence differences between maternal and somatic rRNA variants in zebrafish. rRNA sequence differences between zebrafish maternal and somatic 28S, 18S, 5.8S and 5S rRNAs. Listed are maternal and corresponding somatic rRNA nucleotide positions that differ between the maternal and somatic variants. The table also includes comments based on the analysis of the location and base-pairing status of the specific nucleotides in the 6 hpf zebrafish ribosome structure containing maternal rRNAs.

**Supplementary Table S3:**
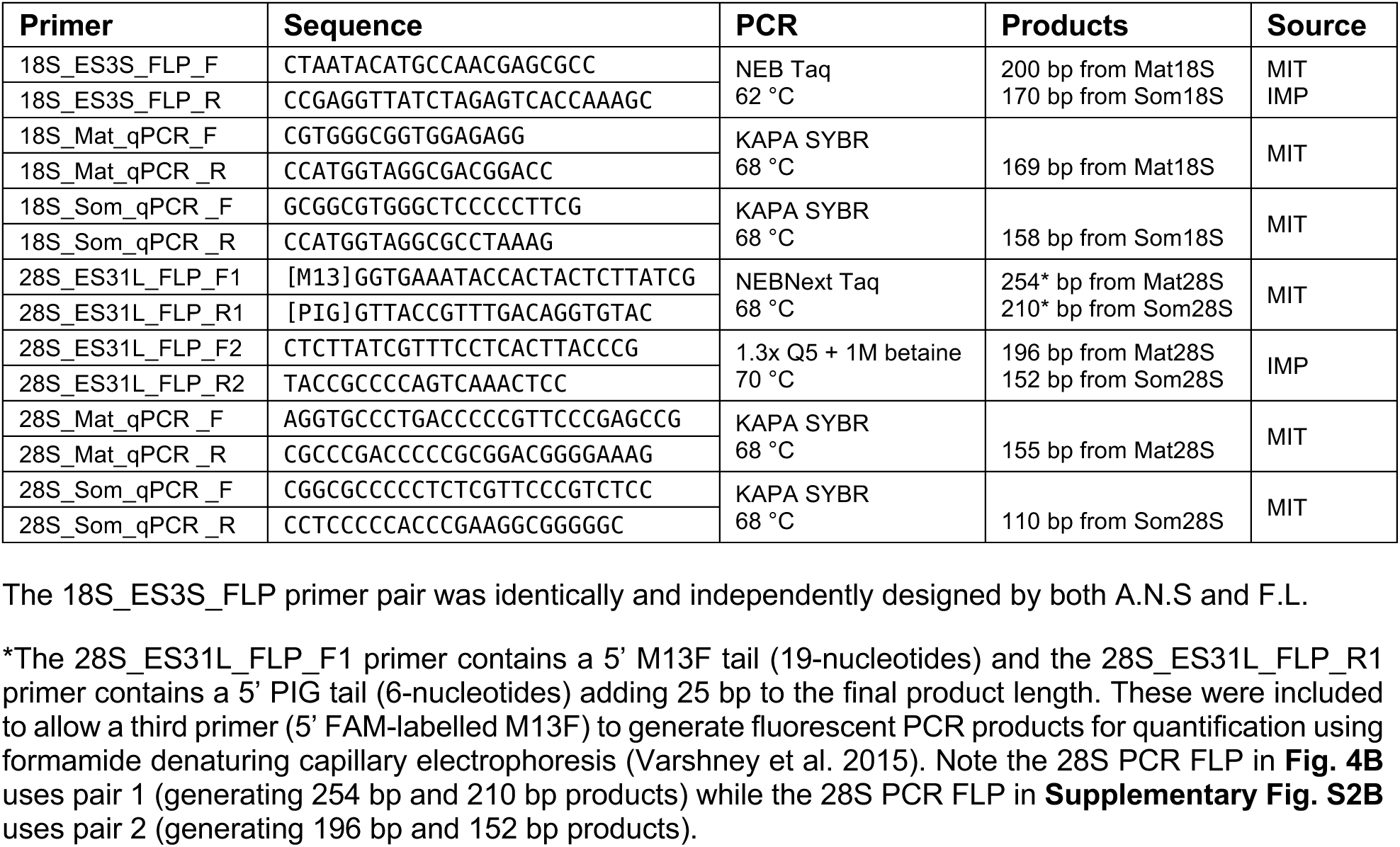
Primer sequences and PCR conditions used to detect maternal and somatic rRNA variants by PCR-based fragment length polymorphism (FLP) and RT-qPCR.

**Movie 1**: Map of the zebrafish maternal ribosome isolated from 6 hpf embryos (PDB: 7OYB; Leesch and Lorenzo-Orts et al., 2023). Maternal rRNA sequences are shown in cartoon representation in shades of yellow (see **Fig. 3A** for the color code); locations of nucleotide differences in zebrafish somatic rRNAs are highlighted in blue (predominantly substitutions), red (predominantly deletions in somatic rRNAs) and purple (substitutions and deletions in somatic rRNAs).

**Movie 2**: Same as Movie 1, but rRNA sequences are shown in surface representation.

**Movie 3**: Same as Movie 2, but cross-section through the ribosome.

## Acknowledgments

We would like to thank Carina Pribitzer for technical assistance; the Mass Spectrometry Facility at the Vienna BioCenter Core Facilities (VBCF) headed by Elisabeth Roitinger and Karl Mechtler, in particular Richard Imre, for the processing and analysis of proteomics data; the IMP aquatics facility personnel, in particular F. Ecker, K. Rattner, J. König and D. Sunjic, for their excellent care of fish, and the Koch Institute Frontier Research Program, the Casey and Family Foundation Cancer Research Fund, the Michael (1957) and Inara Erdei Fund, the Swanson Biotechnology Center Microscopy Core, Grace Phelps from Dr. Lee’s lab at MIT for providing tumor-inducing zebrafish lines; the Koch Institute Zebrafish Core Facility, headed by Adam Amsterdam, and MIT’s BioMicro Center for their support and the VBC RNA Salon and the RNA-Deco-SFB community for providing useful feedback and suggestions. We would also like to thank the entire Pauli and Calo groups for valuable discussions on the project, and for feedback on the manuscript.

## Author Contributions

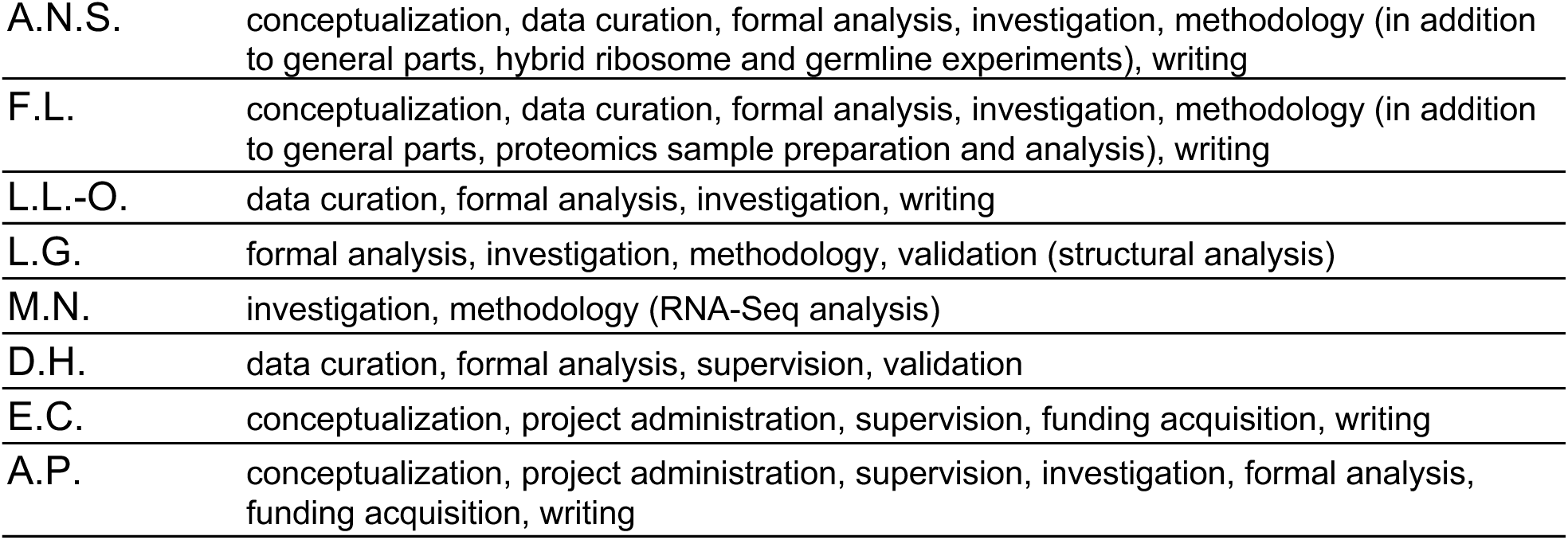

